# Optimizing Xenium In Situ data utility by quality assessment and best practice analysis workflows

**DOI:** 10.1101/2023.02.13.528102

**Authors:** Sergio Marco Salas, Paulo Czarnewski, Louis B. Kuemmerle, Saga Helgadottir, Christoffer Matsson-Langseth, Sebastian Tismeyer, Christophe Avenel, Habib Rehman, Katarina Tiklova, Axel Andersson, Maria Chatzinikolaou, Fabian J. Theis, Malte D. Luecken, Carolina Wählby, Naveed Ishaque, Mats Nilsson

## Abstract

The Xenium In Situ platform is a new spatial transcriptomics product commercialized by 10X Genomics capable of mapping hundreds of transcripts *in situ* at a subcellular resolution. Given the multitude of commercially available spatial transcriptomics technologies, recommendations in choice of platform and analysis guidelines are increasingly important. Herein, we explore eight preview Xenium datasets of the mouse brain and two of human breast cancer by comparing scalability, resolution, data quality, capacities and limitations with eight other spatially resolved transcriptomics technologies. In addition, we benchmarked the performance of multiple open source computational tools when applied to Xenium datasets in tasks including cell segmentation, segmentation-free analysis, selection of spatially variable genes and domain identification, among others. This study serves as the first independent analysis of the performance of Xenium, and provides best-practices and recommendations for analysis of such datasets.

## Introduction

Spatially-resolved transcriptomics (SRT) methods have been crucial for generating molecular atlases in health and disease by placing individual RNA transcripts in the context of tissues^1^. A plethora of SRT methods that leave the tissue intact have been developed in recent years^**2**^. These methods can be divided into different groups. On the one hand, sequencing-based methods (e.g. ST^**3**^, Slide-seq^**4**^) relyon the addition of spatial barcodes to the RNA fragments sequenced later by next generation sequencing, detecting the entire transcriptome with a supracellular (10-100 μm) to subcellular (220 nm) resolution^**5**^. On the other hand, imaging-based methods rely on the detection of individual RNA molecules using fluorescence-based microscopy. Depending on their chemistry, these methods are subdivided into *in situ* hybridization-based (ISH) (e.g. MERFISH^**6**^, SeqFISH^**7**^, EEL^**8**^) and *in situ* sequencing-based techniques (e.g. *in situ* sequencing^**9**^, STARmap^**10**^, SCRINSHOT^**11**^). Despite each technique having specific characteristics^**2**^, imaging-based SRT enables the detection of transcripts at the subcellular level for a selected gene panel over the tissue area. The usefulness of these technologies to investigate the spatial architecture of tissues, propose cell-cell communication events and clarify the roles of different cell types and molecules in disease has led to the emergence of companies interested in commercializing SRT methods. As it occurred with single**-**cell RNA sequencing (scRNA-seq), commercial products based on these techniques promise to facilitate the rapid adoption of methods. One of the most successful examples of these products is Visium, the commercial product of ST by 10X Genomics, which has proven to be a useful tool to study spatial architecture of tissues, despite the limited spatial resolution of the method. Complementing these approaches, several companies have launched imaging-based SRT products (e.g. CosMx by Nanostring, Molecular Cartography by Resolved Biosciences, Esper by Rebus Bioscience, seqFISH by Spatial Genomics). Among those we find Xenium, a 10X Genomics product based on *in situ* sequencing (ISS), that promises to generate maps of hundreds of transcripts at a subcellular resolution. Although Xenium datasets have been used by 10X Genomics to show the potential of the technology^**12**^, an independent exploration and evaluation of the platform is required. In this study, we explore the characteristics, capacities and limitations of data from the Xenium platform. In addition, we explore the performance of several open source computational tools when applied to Xenium data, highlighting the biological insights that they can provide (**Figure 1A**).

**Figure 1:**
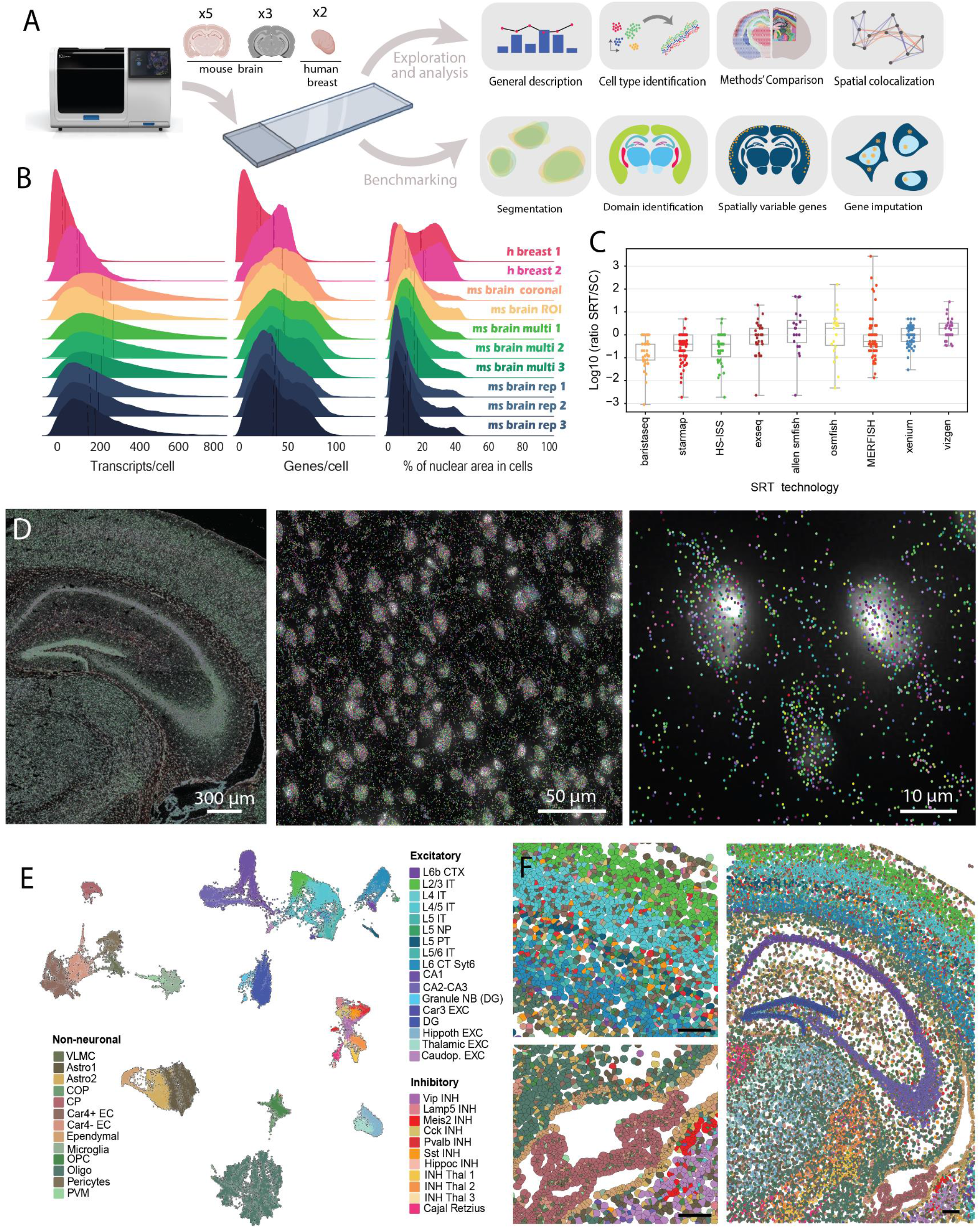
Overview of the analysis and cell type identification using Xenium. **A**. Schematic representation of the overall analysis performed with Xenium datasets. Mouse brain and human breast cancer sections were used to explore the capabilities of Xenium, exploring and benchmarking available methods in several tasks. **B**. Density plot of the transcripts/cell (left), genes/cell (middle) and the percentage of nuclear area (right) identified on each of the datasets. The vertical solid lines indicate the mean and the vertical dashed lines represents the median of each. **C**. Boxplot representing the SRT/scRNA-seq gene efficiency ratios of different SRT methods in log10 scale. Using this scale, a log10 efficiency ratio of 0 indicates the same detection efficiency in both methods. Boxplots represent the distribution of the efficiencies, divided in quartiles, where the central line represents the median efficiency of each technology assessed. Individual gene ratios are represented as individual dots for each method. **D**. Expression maps representing the location of decoded reads in three regions of interest selected from the mouse brain replicate 1 section. Reads are overlaid on top of their corresponding nuclei staining (DAPI) and colored depending on their identity. **E**. UMAP representation of the 41 different cell types identified by Xenium in the 3 adjacent mouse brain regions using unexpanded cells. **F**. Spatial map of the clusters identified in panel E in replicate 1(right) and two regions of interest in the cortex (left, up) and the ependymal / CP (left, down). Cells are represented using the cell boundaries identified by Xenium’s segmentation algorithm after the expansion. Colors correspond to clusters identified in panel E. Scale bar indicate 150 *μ*m.

## Results

### Available Xenium datasets provide high quality measurements of tissue populations

To explore the characteristics of Xenium data, we collected 10 preview datasets generated by 10X Genomics (**Online Methods**) as part of four different experiments. The datasets, obtained with 10X in development chemistry, included (1) five mouse brain coronal sections, obtained in two different experiments, (2) three mouse brain coronal regions of interest (ROI) and (3) two human breast cancer samples which, all together, represented a total of 306.7 million reads and 1.26 million cells (**Extended Data Table 1**). The number of genes profiled (the “panel”) ranged between 210 and 313 genes. All datasets included the 3-dimensional position (x,y,z), gene identity and phred-based quality value (qv) of every decoded read, with 82% (range 77%-91%) of the reads presenting a high quality (qv>20) on average (**Extended Data Table 1**). In addition, cell-segmentation masks and a cell-by-gene matrix containing the expression and position (x,y) of every detected cell were also provided (Online Methods). An average of 229 reads per cell were observed throughout the ten datasets, with 78.5% of the reads being assigned to cells, with no obvious differences between Fresh Frozen (FF) and Formalin-Fixed Paraffin-Embedded (FFPE) sections (**Figure 1B**). Only 0.11% of the cells presented less than 10 assigned reads and were excluded from further analysis, positioning Xenium as a suitable platform to assess cell type frequencies in tissues.

### Xenium elevates ISS gene detection efficiency to rival ISH methods

Mouse brain has been extensively characterized by a wide range of classical and contemporary spatial techniques and for that reason is often used to evaluate technical performance of SRT methods. The cellular composition of the mouse brain has been elucidated through many studies using scRNA-seq^13–17^ and various SRTs^7–9,18–23^, which makes it an excellent tissue for benchmarking SRT methods. We therefore set out to benchmark Xenium against available datasets from the same general region of the mouse brain, primarily isocortex. The imaging-based SRT datasets included the recently published high sensitivity *in situ* sequencing (HS-ISS)^24^, STARmap PLUS^23^, sequential smFISH and MERFISH from the Allen Institute^25^, ExSeq^22^, BaristaSeq^26^, osmFISH^18^ and the Vizgen MERFISH Mouse Brain Receptor Map. For the sequencing-based SRT, we had three different publicly available Visium datasets. Finally, among all the Xenium datasets generated for the mouse brain, we used the 3 full coronal sections, named as “multisection” in Figure 1B, as they represented the most updated version of Xenium’s chemistry. Anatomical regions were manually annotated in the different SRT datasets to guarantee a fair comparison between similar regions (**Online methods, Extended Data Figure 1**) and then proceeded to compare the cells from annotated regions from the cortex. One inherent feature of the imaging-based SRT technologies is that they are targeted, which means that the number of detected molecules per cell varies depending on the probes used. We therefore calculated the detection efficiency of individual genes measured by each technology by comparing them to a reference scRNA-seq dataset^17^. Overall, we observed Xenium to be more sensitive than all other ISS-based techniques, presenting a similar sensitivity to some ISH-based technologies (**Figure 1C**). In addition, Xenium presented an overall detection efficiency similar to scRNA-seq (Chromium v2) (mean ratio=1.07). However, due to the nature of the method, the efficiency ratio of individual genes presented a high variability, ranging from 0.21 to 2.0. A negative correlation (r=0.53) was found between the level of expression in scRNA-seq and the SRT/scRNA-seq gene efficiency ratios (**Extended Data Figure 2A**), indicating that lower expressors from scRNA-seq are detected with a high efficiency *in situ* and *vice versa*. This could be explained by the different number of padlock probes added for each gene, which was done to tune the detection efficiency of each gene in a customized way in order to reduce optical crowding limitations (**Extended Data Figure 2B**). However, the variability between genes targeted with the same number of probes suggest that additional factors play a role in determining the SRT/scRNA-seq gene efficiency ratios.

Finally, we compared Xenium against Visium^3^, the most widely used spatial transcriptomics platform. Since Visium doesn’t have single cell resolution, cortical and hippocampal regions were subset from both datasets, comparing the pseudo bulk reads detected by each method for each gene. Xenium was found to be much more efficient than Visium on a tissue level, with a median of 21 times more reads detected using Xenium compared to Visium Fresh Frozen (FF) (**Extended Data Figure 2C**).

### Identified populations are influenced by the expansion of nuclei

For further exploring Xenium datasets’ characteristics, we focused on the three adjacent tissue slices imaged in the mouse cortex and hippocampal regions (**Figure 1D**). These sections exhibited a high similarity in terms of read quality and counts (**Extended Data Figure 2D**) (Pearson’s r = 0.997), which allowed us to use them as technical replicates throughout the study. Xenium’s customized cell identification algorithm consists of an initial nuclei segmentation based on DAPI, followed by an expansion of the segmentation masks. Using the cell-by-gene matrix of segmented nuclei, we identified a total of 41 cell types in the three mouse brain datasets that were consistent with those previously identified in scRNA-seq experiments (**Figure 1E, Extended Data Figure 2E**) (Online Methods), demonstrating that nuclei segmentation masks contain sufficient information to decipher the identity of individual cells *in situ*. These cell types were then mapped back onto the tissue to create a cell-type map (**Figure 1F**). The proportion of detected cell types was highly similar across all the three adjacent slices (Pearson’s r = 0.99) (**Extended Data Figure 2 D-F**). When assigning these annotated cells to anatomical tissue domains (Online Methods) we observed a consistent distribution of domain-specific cell types^13^(**Extended Data Figure 2 G-H**).

Despite the transcripts assigned to cells based on nuclear segmentation being sufficient to define cell populations, some cytoplasmic reads might be lost in the process. Under this assumption, Xenium’s nuclear segmentation is followed by an expansion of 15 μm. When identifying cell types using this expanded cell-by-gene matrix, many of the cell types were divided into region-specific clusters, in contrast with the homogeneous cell types identified using unexpanded segmentation (**Extended Data Figure 3 A,B**). In addition, clusters in the UMAP based on expanded nuclei were globally embedded by region (**Extended Data Figure 3C**). For instance, oligodendrocytes in the thalamus were grouped together with astrocytes in the thalamus, rather than with other oligodendrocytes (**Extended Data Figure 3 D-E**). This indicates that expansion captures domain-specific expression signatures, influencing the clustering. Furthermore, the clusters detected didn’t present clear marker genes and, compared with nuclei-based clusters, some biologically relevant cell types were lost in favor of region-specific clusters (**Extended Data Figure 3 F-G**).

To better understand the region specific effect of expansion, which we suspected was due to misassignment of spots from neighboring cells, we defined nuclear expression signatures for each of the identified cell types, as well as background expression signatures for each of the manually annotated domains. Taking these signatures as a reference, we found that including transcripts beyond 8.24 um from the cell centroid resulted in a higher gene expression correlation to background domain-specific expression signatures compared to nuclear cell type-specific expression signatures (**Figure 2A-C, Extended Data Figure 4 A-B**). This distance likely represents the average radius of identified cells in the tissue, including both nuclei and cytoplasm. Since, on average, nuclei in this dataset presented a radius of 4.85 μm, the ideal expansion of cells in the samples analyzed should be of 3.38 μm. Different cell types presented, though, different average distances, representing their diverse cellular morphologies. As a consequence, segmentation strategies based on the identification of nuclei followed by a rigid expansion might not be the optimal solution.

**Figure 2.**
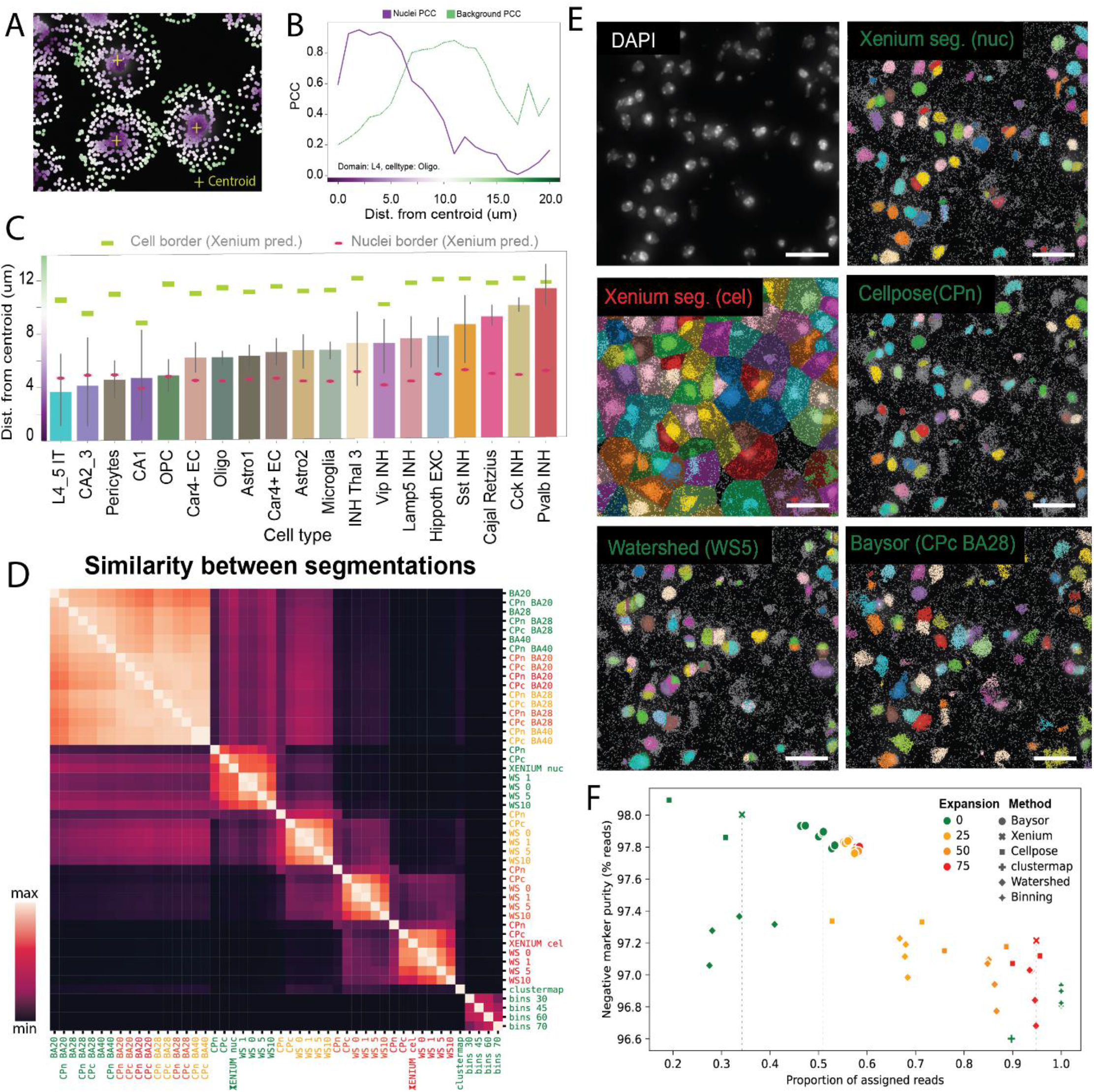
Exploring segmentation in Xenium. **A**. Region of interest of one of the mouse brain sections. Reads are overlaid on a DAPI staining and colored depending the distance to their closest cell centroid. Color legend is present in Figure 2B. **B**. Pearson correlation coefficient (PCC) of OPC to the nuclear OPC mean expression signature in Layer 4 of the cortex (purple) and the PCC to the background Layer 6a-specific expression signature (green, dashed) depending of the distance to the cell centroid. **C**. Bar plot indicating distance of the intersection between nuclei and the domain-specific background as represented at Figure 2B for a subset of the clusters identified at Figure 1E. The intersection point is calculated for all the possible combinations of cell types and domains, provided that the assessed cell type has at least 5000 reads assigned to the assessed domain. Error bars represent the 95% confidence interval. The mean nuclei and cell radius of every cluster are represented using red and green rectangles respectively. **D**. Adjusted rand index (ARI) between the different outputs produced by combinations of segmentation algorithms, hyperparameters and expansions. Segmentation methods included Cellpose (CPn: nuclei, CPc: cyto models), binning (bins), clustermap, watershed (WA), Baysor (BA) and cellpose combined with Baysor (CPc BA/CPn BA).Xenium segmentation were also included in the comparison (XENIUM cel, XENIUM nuc). Hyperparameters for each method are described in methods. Methods on the y-axis were colored depending on the expansion performed after segmentation. Scale bars represent 25 μm **E**. Cells identified using different segmentation algorithms in a region of interest outlined in Extended Data 4C corresponding to L6. DAPI is represented as a background and color-specific masks represent individual cells. Segmentation strategies represent different segmentation outputs, as described in Figure 2E. **F**. Scatter plot representing the number of reads assigned (x-axis) and the negative marker purity (y-axis) of different assessed segmentation strategies. The name and color of each segmentation strategy are represented as in Figure 2E.

### Standard Xenium segmentation is outperformed by Baysor and Cellpose

The influence of cell segmentation on cell-typing accuracy motivated us to explore alternative segmentation methods. For this, we benchmarked the performance of the segmentation provided by Xenium against commonly used segmentation strategies (Online Methods). These strategies can be broadly classified as: staining-based, if the position of cells is determined by an auxiliary staining like DAPI (watershed^27^, Cellpose^28^); read-based, where cells are defined based on the read density and composition of tissues (Baysor^29^); or mixed models, where both staining and the position of reads is used for defining cells (Baysor^29^, Clustermap^30^). Segmentation based on equally distributed bins across the tissue (binning) was also included in the comparison as an example of simplistic segmentation. In addition, different cell expansions were applied on top of each segmentation output (0, 5, 11, 16 μm).

We estimated the similarity between segmentation strategies using Adjusted Rand Index (ARI), identifying groups of strategies performing similarly (**Figure 2 D-E, Extended Data Figure 4 C-D**). Staining-based strategies applied to DAPI generated similar outputs, with cell expansion as the driving force of their differences. In addition, Baysor-based, Clustermap-based and binning strategies were found to cluster by method, indicating distinct method-specific segmentation output. Alongside quantitative assessment of segmentation similarity, we also employed quantitative metrics to evaluate the performance of segmentation strategies. We defined the best segmentation strategy as the one maximizing the proportion of reads assigned to cells while maintaining specific expression patterns, quantified by Negative Marker Purity (NMP) (Online Methods) (**Figure 2F, Extended Data Figure 5**). NMP measures, based on a reference scRNA-seq^17^, the percentage of reads detected in cell types that are expected to express the read’s gene. We identified Baysor-based strategies and specifically Baysor combined with a Cellpose-based segmentation (BA28-CPc) as best segmentation strategy (**Figure 2F**). Although no additional cell types were detected using BA28-CPc segmentation in comparison with Xenium’s nuclear segmentation (**Extended Data Figure 4E**), we observed a median increase of 44 reads per cell (127 vs 83).

### Segmentation-free models provide an alternative to classical workflows

Segmentation-free models, where the identification of spatially-resolved molecular signatures is done independently from the segmentation of individual cells have proven to be an alternative approach to the traditional analytical workflow. The more parsimonious nature of segmentation-free approaches, which avoid modeling cellular structures and operate on the immediately detected signal instead allows for a different type of analysis. First, they can be used to investigate local signal properties and fidelity before the spatial accumulation into cells. With the aim of exploring cellular and subcellular patterns, we applied two of these approaches, SSAM^31^ and Points2Regions^32^. Using SSAM *de novo mode*, a total of 26 cell type-specific clusters were identified. The comparison between these clusters and the cell types identified in Figure 1E highlights the capacity of these methods to capture cell type-specific expression patterns (**Figure 3A-C**). Since, unlike in segmentation-based methods, all the reads are included in the analysis, extranuclear reads were consistently assigned to specific signatures without the necessity of performing segmentation. Moreover, the relatively simple and resource-effective SSAM algorithm can incorporate information of molecule placement along the z-axis and potentially identify subcellular structures and compartments. We used Xenium’s 3-dimensional coordinates to detect cases of possible mixed-source signals originating from cells overlapping in the z-dimension (**Figure 3D-E**). The presence of these cases (**Extended Data Figure 6A**)highlights the importance of considering spatial datasets as 3-dimensional maps, rather than simplifying their position to 2-dimensional coordinates.

**Figure 3.**
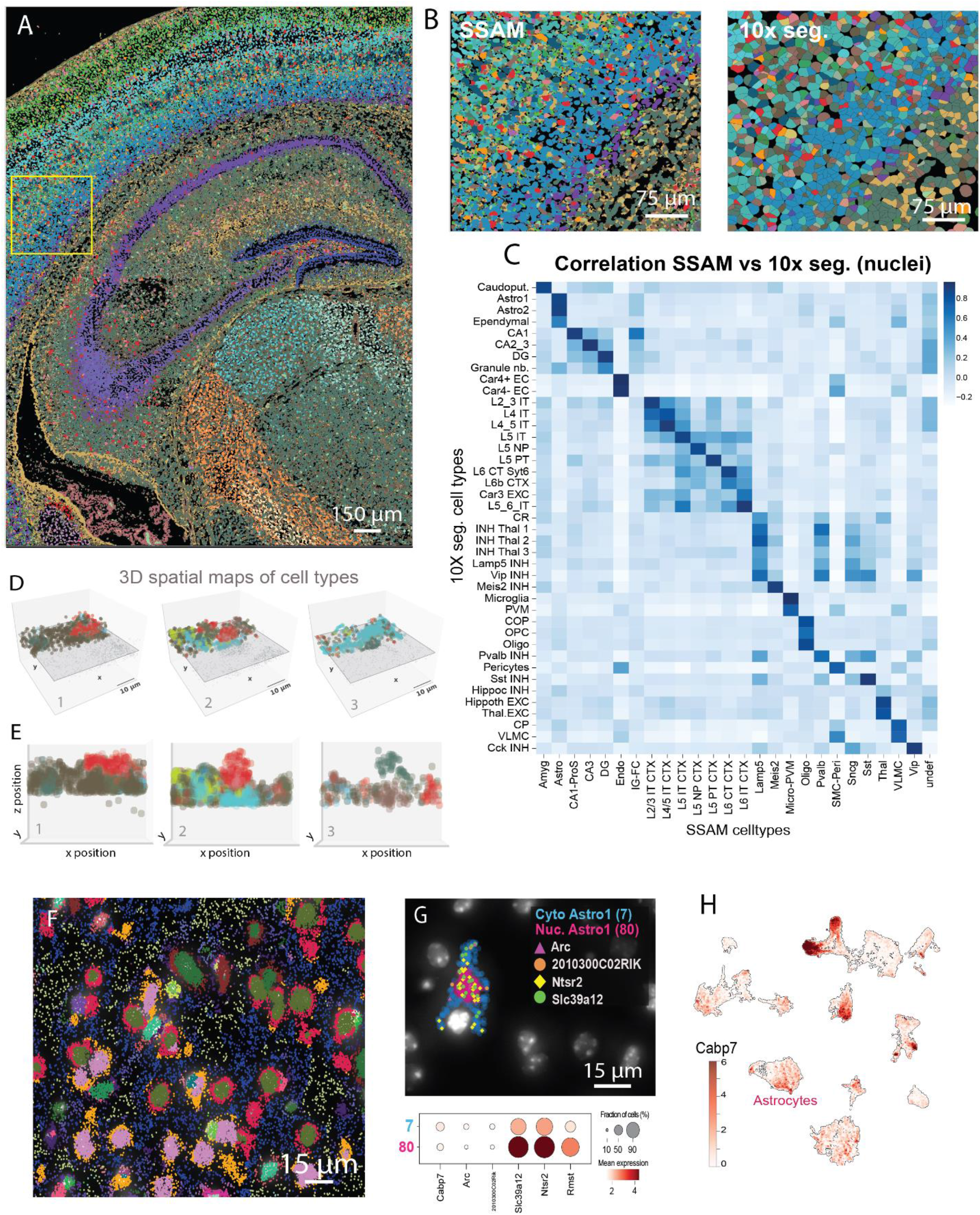
Segmentation-free analyses highlight 3D and subcellular patterns. **A**. Cell type map of the signatures identified in one of the mouse brain adjacent slides (slide 2). Colors represent the different cell types identified, using the same color map as in Figure 1D. A yellow square represents the region of interest represented in Figure 3B. **B**. Representation of cell types using SSAM (left) and the default 10x segmentation (nuclear segmentation, right) in the cortical region of interest highlighted in Figure 3A. **C**. Pearson correlation between the cell type-specific signatures described using SSAM and those found in Figure 1E using nuclei-based segmentation. **D-E**. Spatial representation in 3D (D) and 2D (E) of overlapping cells found in the mouse brain section analyzed in Figure 3A. **F**. Representation of the subcellular clusters identified by Points2Regions in the mouse brain section analyzed in Figure 3A. Individual spots are overlaid on top of a DAPI staining and colored by the assigned cluster. **G**. Representation of astrocyte-associated clusters with different subcellular localization (up). The reads assigned to some genes with a subcellular patterning in astrocytes are represented also in the map (up). The expression of the top 3 differentially expressed genes between subcellular astrocytic clusters are represented in the form of a dot plot (bottom). **H**. UMAP representing the expression of Cabp7 in the different cell types identified in Figure 1E. Astrocytes are highlighted.

Besides identifying cell types, we explored whether the read density of Xenium datasets could be sufficient to identify subcellular expression patterns. Using Points2Regions, we identified a total of 100 clusters representing subcellular patterns (**Figure 3F**). By coupling them with the output of Xenium’s nuclear segmentation, these clusters were classified in nuclear, cytoplasmic or extracellular. Furthermore, many of these clusters could be associated with specific cell types. We identified subtle yet specific expression differences between nuclear and cytoplasmic clusters associated with the same population (**Figure 3 G-H**), suggesting that Xenium’s signal density enables the identification of subcellular structures *in situ*.

### Benchmarking the performance of gene imputation tools on Xenium datasets

Targeted SRT methods are generally limited by the number of genes measured simultaneously. Imputation approaches overcome this by predicting gene expression from a reference scRNA-seq onto the cellular-resolution SRT dataset^33^. We sought out to benchmark the performance of five methods (gimVI^34^, SpaGE^35^, Tangram^10^, SpaOTsc^36^ and NovoSpaRc^37^) using the workflow designed by Li et al.^33^ and taking as a reference a cortical and hippocampal atlas^17^. Brain regions not included in the reference scRNA-seq were excluded from comparison (**Figure 4A, Extended Data Figure 6B**) (Online Methods).

**Figure 4.**
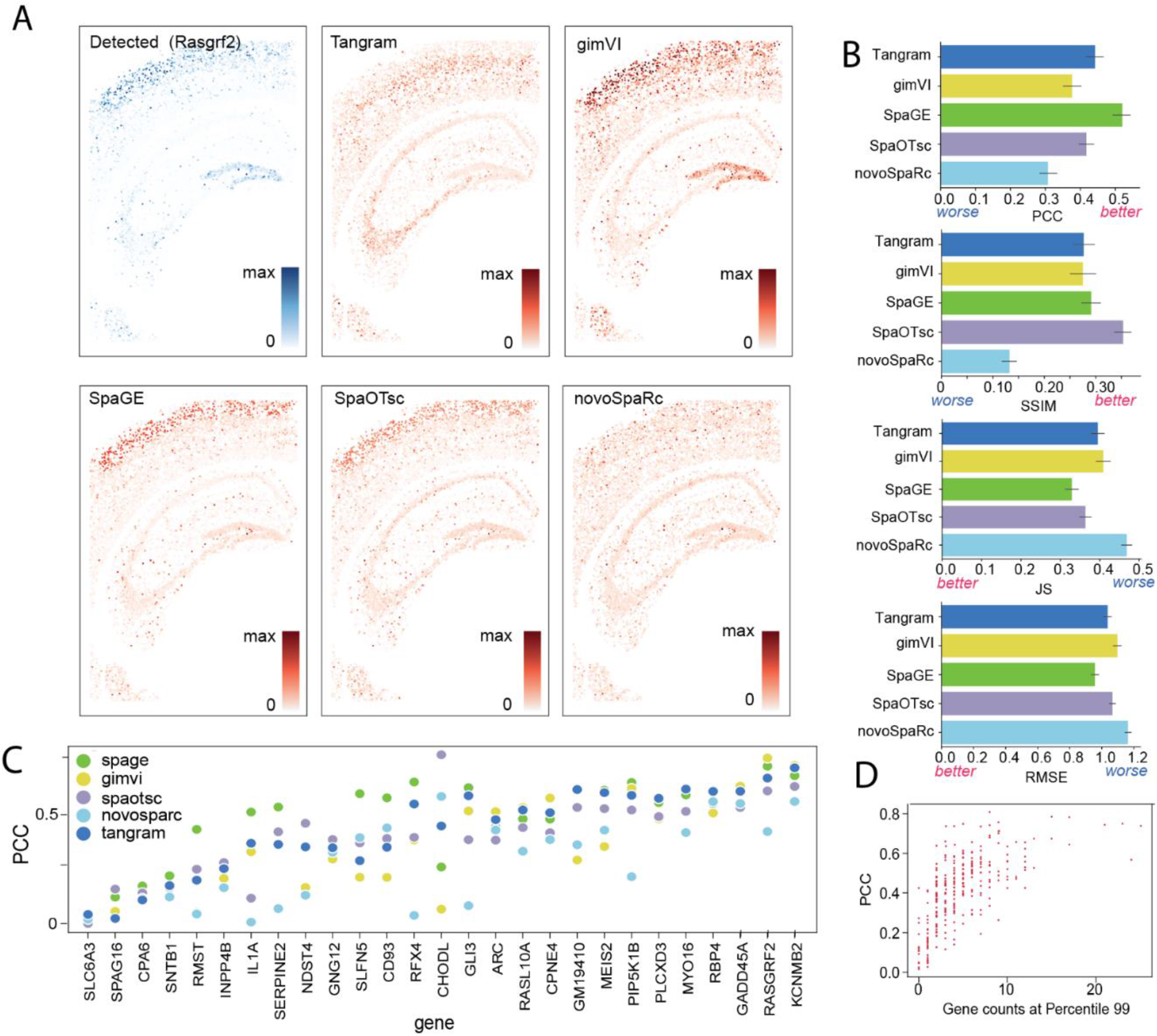
Benchmarking imputation algorithms combined with Xenium. **A**. Detected and imputed expression of the gene Rasgrf2 in the cortex and hippocampus using different imputation algorithms. **B**. Performance of different gene imputation methods as described by 4 different metrics: pearson correlation coefficient (PCC), structural similarity index (SSIM), the Jensen-Shannon divergence (JS) and root mean square error (RMSE). Data are presented as mean values ± 95% confidence intervals. **C**. Scatter plot representing the PCC found between the detected expression and the imputed expression for specific genes using different gene imputation algorithms. **D**. Relation between the imputation accuracy scores (as PCC) of individual genes and the detected gene counts of the genes at percentile 99, represented as a scatter plot.

Imputation performance was assessed by Pearson correlation coefficient (PCC), structural similarity index (SSIM), root mean square error (RMSE) and the Jensen-Shannon divergence (JS), where a higher PCC/SSIM and a lower RMSE/JS value indicates better prediction accuracy. Using these metrics, we consistently identified SpaGE as the best performing method (**Figure 4B**). In addition, Tangram and SpaOTsc achieved an overall high performance. Surprisingly, gimVI performance was found to be lower than the one reported when using it to integrate scRNA-seq with other SRT technologies^33^.

Since the workflow used relies on comparing detected and imputed gene expression of individual genes, we conceived this as a good way to identify genes with an overall low agreement between scRNA-seq and Xenium. By quantifying these gene-specific differences using PCC across all genes and methods, we observed an enormous difference in the imputation performance between methods. These differences were consistent across imputation methods (**Figure 4C**) and were correlated with the level of expression of the genes *in situ* (**Figure 4D**) (PCC=0.63).

### Assessing computational tools to explore tissue architecture

Imaging-based SRT has the ability to recover the spatial location of individual cells which can be used to decipher the organization of tissues’ intrinsic architecture. By utilizing computational frameworks for the exploration of spatial datasets implemented in Squidpy^38^ and Giotto^39^, we identified the major spatial regions in mouse brain (**Figure. 5A-B**). While some cell types like excitatory neurons in the hippocampus presented a more restricted connectivity due to their specific location, other cell types such as astrocytes or OPCs presented a broader number of connections to other cell types, in line with their known broad distribution.

**Figure 5.**
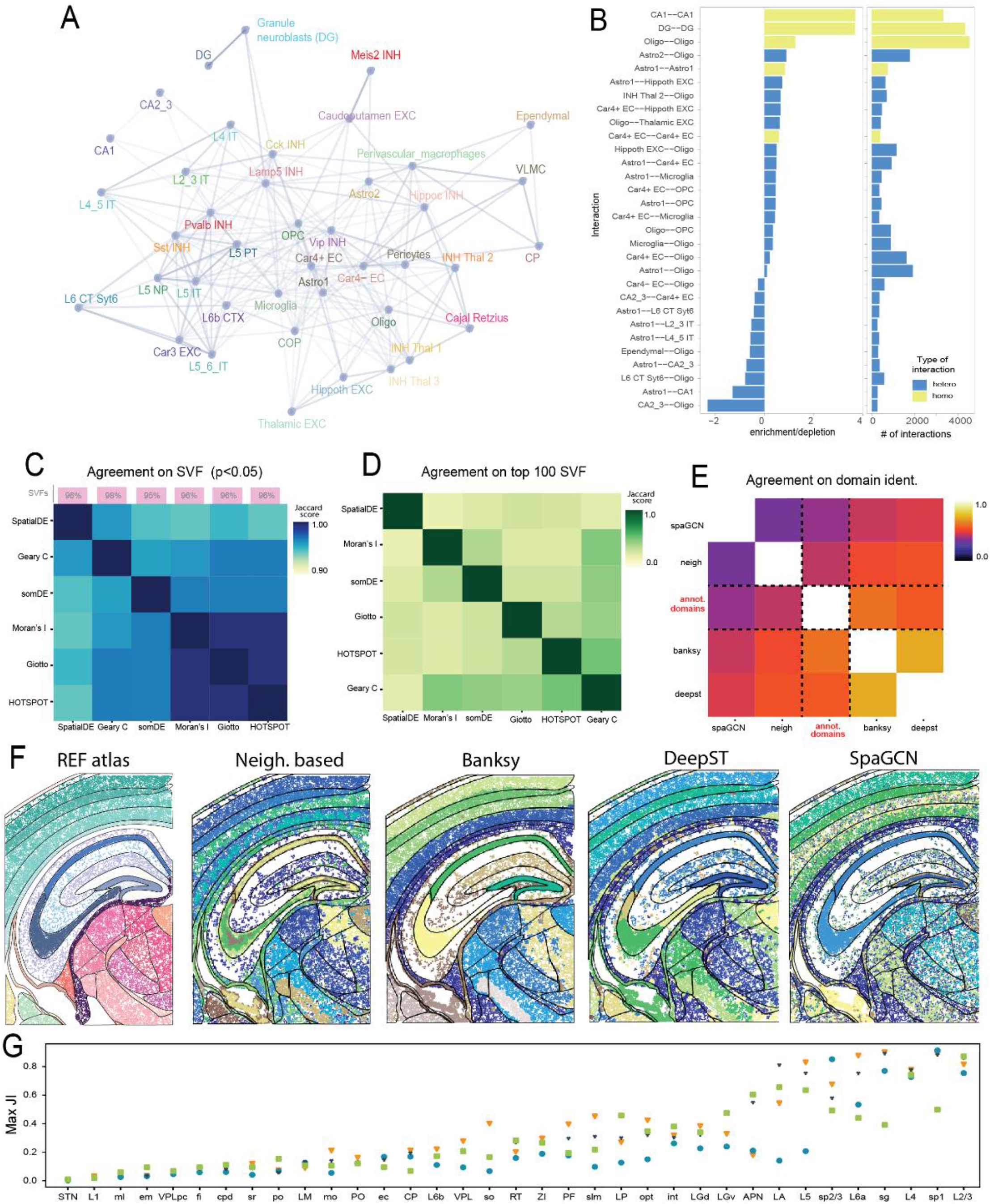
Spatial characterization using Xenium. **A**. Network representing the spatial enrichment between different cell types. Nodes represent each cell type annotated in Figure 1C, with colors corresponding to the ones used in the mentioned panel. Edges between cell types presenting a positive significant enrichment (p>0.05) are drawn. Edges’ width corresponds to the enrichment scores. **B**. Bar plot representing the enrichment/repletion score (left) and number of connections (right) between different cell type pairs. Only pairs presenting a p-value>0.005 and a total of interactions larger than 250 are plotted. Bars are colored depending on the type of interaction. **C**. Heatmap representing the Jaccard score obtained by comparing the significant spatially variable features (SVF) (p<0.05) detected by different algorithms. Small bar plots on the top of each column represent the percentage of genes identified as SVF by each method. **D**. Heatmap representing the agreement between different SVF methods when selecting the top 100 SVF by each method. **E**. Heatmap representing the agreement between different algorithms in identifying tissue domains in mouse brain sections. **F**. Spatial map of the manually annotated domains identified in the mouse brain section (replicate 1, left) and the domains identified by different algorithms including (Banksy, DeepST, SpaGCN and neighborhood-based region identification) **G**. Scatter plot representing the maximum Jaccard Index (JI) (y-axis) between each reference-based domain (x-axis) and the different domains predicted by the different domain finder algorithms.

Despite the architecture of cell types, the identification of spatially variable features (SVF) is useful to discriminate genes explaining the main spatial variation patterns within the tissue. The plethora of methods available for this task, though, could lead to slightly different conclusions depending on the algorithm used. When comparing the most commonly used metrics and algorithms developed for this task (Moran’s I^38^, Geary’s C^38^, Hotspot^40^, SomDE^41^, SpatialDE^42^, Giotto^39^) we observed that a mean of 96% of the genes included in the Xenium’s panels were considered as SVF (pval<0.05) (**Figure 5C**). All methods presented a high consistency in identifying SVFs, with overall Jaccard scores above 0.95, with the only exception of SpatialDE. The consistency of different algorithms in identifying SVF is lost when comparing the most variable genes identified by every method (**Figure 5D**). As a consequence, a selection based on significance would be preferred in contrast with the selection of the top SVFs. Since most of the genes targeted are consistently found to be spatially variable, though, a gene selection of SVF in Xenium would not be as useful as in other untargeted spatial transcriptomics technologies.

Identifying tissue architecture can be helpful to understand its function. Identifying the most reliable tools to define these domains is of general interest, although no independent comparison is yet available. Therefore, we benchmarked 4 domain finder algorithms (neighborhood-based^43^, Banksy^44^, DeepST^45^, SpaGCN) against the regions identified by expert manual annotation using the coronal P56 section from Allen Brain Atlas^46^ (**Figure 5D-E**). A total of 36 domains were manually annotated, and thus each method was adjusted to predict a similar number of domains (35-37). We found the domains predicted by Banksy to be the most similar ones compared to the manual annotation (Adjusted Rand Index = 0.59) (**Figure 5D**). Nevertheless, there were consistent differences in the identification of specific domains (**Figure 5F**). While cortical and hippocampal regions were overall accurately predicted, others like thalamic regions were consistently misidentified (**Figure 5F**). As expected, expression differences between domains were correlated with a good identification of domains, being especially the case in most transcriptionally distinct domains.

### Best practices for processing and analyzing Xenium datasets

As seen previously, the way Xenium data is processed and analyzed can distorsion the biological significance of the results obtained. For this, and based on the evidences shown through the manuscript, we propose an optimal processing and analysis of Xenium datasets. We have condensed this information in an end-to-end pipeline (**Code Availability**) with the aim of helping Xenium users to take the most out of their data.

Briefly, taking the data obtained from Xenium as an input, the first key step for an optimal processing of the data is cellular segmentation. The optimal algorithm for this consist on two steps: (1) the identification nuclei using Cellpose^28^ and (2) the assignment of reads to individual cells using Baysor^29^. Cellular expansion is not needed, since extranuclear reads are assigned using Baysor. If cellular segmentation results in a poor performance, segmentation free methods like SSAM^31^ or Points2Regions^32^ can be used to identify molecular signatures without the necessity of identifying individual cells. After segmenting individual cells, a cell-by-gene matrix is obtained. Taking this as an input, cell type identification can be achieved using standard scRNA-seq workflows consisting of (1) cell filtering, (2) log-transformation and normalization, (3) identification of the main PCs, (4) low-dimensional reduction and (5) clustering. The selection of SVF was found to be an unecessary step in this process due to the targeted nature of the method. In addition, if sc/snRNA-seq is available, gene imputation can be performed using algorithms such as SpaGE^35^or Tangram^10^. Finally, for the identification of domains, using Banksy can be a powerful solution.

## Discussion

In this study we presented an independent exploration and evaluation of Xenium *in situ*datasets. The assessed technology can be used to generate highly multiplexed spatial maps of RNA molecules at a subcellular resolution. The number of reads per cell identified, in the range of hundreds, and the area of the tissues characterized facilitate the easy identification of cellular populations *in situ*.

The Xenium *in situ* platform showed a detection efficiency comparable to Chromium v2, representing a remarkable increase in sensitivity in comparison with other ISS-based methods. Among SRT methods, however, these comparisons need to be taken with caution, as gene-specific sensitivities depend on the probes used to target them. RCA-based technologies, like Xenium, enable the tuning of molecules detected per gene. We believe that modifying these levels for intentionally decreasing the detection levels of high expressors or, more importantly, increasing the detection levels of lowly expressed genes is one of the main advantages of this approach in comparison with other commercial solutions. Although Xenium presents a considerably higher detection efficiency than Visium, when compared with some other available ISH based SRT products and platforms, its mean detection efficiency was found to be slightly lower. Despite this being one of the most important features of SRT methods, further independent comparison of technical aspects such as the imaging time, cost of individual experiments, simplicity of the technique and reproducibility would be needed to be more comprehensive compared between the different technologies and commercial solutions.

Several processing steps were identified to be crucial when analyzing Xenium datasets with segmentation being highlighted as one of the most important. With the current segmentation provided by 10X Genomics, cells’ nuclei were overall nicely identified and their information found to be sufficient for identifying the main cell populations present in the sections analyzed. We identified that the expansion of cells after segmentation can have detrimental effects in the characterization of cell populations due to the assignment of domain-specific reads to cells from neighboring cells. By comparing alternative segmentation strategies to define cell populations, we identified Baysor combined with Cellpose segmentation to outperform the rest of the strategies, being capable to define individual cells based both on the density and identity of individual reads *in situ*. This strategy allows Baysor to identify cells with different sizes, composition and to even detect cells that didn’t present their nuclei in the analyzed section due to sectioning. Despite its overall good performance, Baysor struggled in identifying cells in dense homogenous areas such as the hippocampus. Overlapping cells were identified as a recurrent yet difficult to segment case due to the 2-dimensional nature of most of the algorithms used. Thus, the implementation of new segmentation algorithms that consider the 3-dimensional structure of the data and the inclusion of additional stainings for cellular membranes would facilitate the correct identification of individual cells.

In this direction, segmentation-free cell typing methods represent an alternative to the typical segmentation-then-clustering workflow, being able to identify cell populations and subcellular patterns. The exploration of subcellular patterns represents an underexplored layer of information that is now becoming accessible due to the sensitivity and resolution of new imaging-based SRT methods such as Xenium. The combination of these technologies with systematic analytical approaches using segmentation-free algorithms or tools like Bento^47^ could facilitate the understanding of RNA biology at a subcellular level.

Due to the vast availability of methods, just a subset of the most popular methods were included in the benchmarking of the different tasks and thus, for each task the top performer method found might not be the best available method. In addition, some of the published algorithms approached the tasks in an alternative manner, resulting in non-comparable outputs (i.e. SpaGCN^48^ identifies domain-specific SVFs) and were not suitable for benchmarking. Overall, despite some efforts being made, a more systematic benchmarking of these and other algorithms considering datasets generated in different tissues with different experimental designs would be beneficial to decipher when to use each algorithm. For this, the accessibility to new and more diverse datasets would be recommended.

All in all, Xenium represents an overall improvement in comparison with other RCA-based technologies. Its increased detection efficiency, together with its resolution enables the identification of cell types in space, being a useful tool to explore spatial biology. Furthermore, it can be easily complemented by open source algorithms, which further expand the analytical possibilities of these datasets.

## Online Methods

### Xenium datasets processing

The 10 datasets included in this study were formatted in the *anndata* format using a customized function (https://github.com/Moldia/Xenium_benchmarking)and processed using Scanpy (v1.9.1.). Cell-by-gene matrices provided by 10X genomics were log-transformed and normalized. In order to identify clusters, neighborhood graphs were computed considering 40 principal components and 12 neighbors, followed by Leiden clustering. Cell-type annotations were performed using differentially expressed marker genes previously described in Yao et. al.^17^. The same preprocessing steps were performed on cell-by-gene matrices obtained for alternative segmentation methods.

### Annotation of mouse brain architecture

Tissue domains from SRT datasets were manually annotated using the mouse coronal P56 sample from Allen Brain Atlas^46^. Datasets were first processed as described above and then used to generate plots containing cell cluster or spot identity, overlaid on their respective DAPI-, immunofluorescence- or H&E-stained images. Region delimitations were created using enclosed vectorized Bézier-curves, which could then be saved as polygons in scalable vector graphics format (.svg). These annotations were then integrated to the cell and/or spot coordinate system from each sample, respectively, allowing both classification of cells and/or spots and projection of annotated regions onto the datasets for inspection and downstream analysis (see below). This approach promotes a more reproducible annotation of regions of interest, which can be further modified or updated as necessary, while also being scalable and machine readable. All annotations and detailed instructions on how to use them are freely available at https://github.com/Moldia/Xenium_benchmarking.

### Single-cell RNA sequencing processing

A subset of the scRNA-seq dataset from Yao et. al.^17^ was used in this study. For generating this subset, guaranteeing the equal representation of all populations, populations were subsampled to a maximum of 500 cells each. The resulting subset consisted of 150K cells further used for integration and annotation of the populations found *in situ*. Cell type signatures of each cell type were also obtained for the different levels of annotations presented in Yao et. al. These signatures were used as an input in several cell typing methods, such as SSAM.

### Benchmarking Xenium against other SRT methods

For comparing the gene detection efficiency between different SRT methods, only cells included within the manually annotated isocortex regions present in all image-based datasets were used (L2/3,L4,L5,L6b).In the side by side comparison between Xenium and Visium, manually annotated regions present in both datasets were considered. Cell-by-gene matrices from all available datasets were used for comparison. For comparison with scRNA-seq, a subset of the scRNA-seq dataset from Yao et. al.^17^ was considered (see previous section). Raw counts were compared between all methods. Since SRT methods measure a limited panel of genes, there wasn’t any gene detected in all experiments. Genes present in at least 4 datasets were further used for comparison. Taking scRNA-seq as the reference dataset, the detection efficiency of each selected gene for each method was assessed by (1) identifying positive cells for a certain gene in both methods (cells with more than 1 read), (2) computing the median expression of a gene in the population of positive cells and (3) computing the ratio between the two medians. Since we aimed to identify the efficiency of each specific gene in each SRT method, we divided the SRT median by the scRNA-seq median. We considered solely positive cells for the estimation in order to reduce the effect of having slightly different cell compositions in the different datasets.

### Benchmarking segmentation algorithms

We benchmarked the segmentation methods Baysor, Watershed, Cellpose, and Clustermap in comparison with segmentations provided by 10x and equally distributed bins across the tissue (binning) with a custom pipeline. All segmentation masks were expanded with 0, 5, 11, and 16 μm (0 - 75 pixels). Watershed was applied using squidpy (v1.2.3) with Gaussian blurring of standard deviations 0, 1, 5, and 10 pixels (determining the naming WS0 - WS10) and Otsu’s thresholding. Baysor (v0.5.2) was tested with scale parameters 20, 28, 40 pixels (BA20 - BA40) and optionally a prior segmentation with default prior-segmentation-confidence of 0.2 (e.g. CPc BA28). The Cellpose (v2.0.5) deep learning models “nuclei” (CPn) and “cyto” (CPc) were applied on the DAPI channel in their vanilla versions. Clustermap (github code from Nov 8, 22) was applied with gaussian blurring, xy_radius 15, and cell_num_threshold of 0.0001.

The pipeline takes as input the DAPI image, the gene spots with cell type annotations and x and y coordinates, as well as a scRNA-seq reference dataset with cell type annotations that are matched with the cell type naming for the spatial cells. The dataset described in methods “scRNA-seq processing” was used as reference. The cell type annotations on the nuclear segmentation from 10X (methods “Xenium datasets processing”) was provided with the spots. After each segmentation run a count per gene matrix of the cells were generated and cell types were assigned. As new cells are identified with every segmentation run, cell types of the new cells were assigned based on a spots’ majority vote of the previous nuclear segmentation-based annotation. For each segmentation output the pipeline measured the proportion of assigned reads, number of identified cells, the median and 5th percentile of reads per cell, and the median and 5th percentile of genes per cell. Further we introduced and measured the Negative Marker Purity (NMP) metric which is based on the assignment distribution of reads from negative markers: A negative marker for a set of “negative cell types” is defined, based on the single cell reference, as a gene that is expressed in less than 0.5% of cells in each of the negative cell types. Two versions for the metric were defined: 1. (NMP reads) the percentage of reads of negative markers in the cell types expected to express the gene, and 2. (NMP cells) the percentage of positive cells of cell types that are expected to be positive for a given gene relative to all measured positive cells for the gene.

### Benchmarking gene imputation algorithms

We evaluated the performance of five integration methods (gimVI, SpaGE, Tangram, SpaOTsc and NovoSpaRc) following the published Jupyter notebook by Li et al^33^. (https://github.com/QuKunLab/SpatialBenchmarking/blob/main/BLAST_GenePrediction.ipynb). We used the raw normalized expression matrices for both the scRNA-seq and spatial transcriptomics datasets for input of the integration methods. Since the spatial transcriptomics dataset contained less than 100 detected genes, we built the ground truth of our dataset using genes detected in both the spatial transcriptomics and scRNA-seq datasets (total 284 genes). For the evaluation we used tenfold cross validation, where we for each iteration divided the genes into 10 portions, 9 of which were used for training and one for prediction. Because of the large sizes of our datasets (scRNA-seq dataset ~150,000 cells, spatial transcriptomics dataset ~80,000 spots) we downsampled the datasets by a factor of ten, the lower amount of computational resources consumed by the integration methods not compromising the results.

### Benchmarking domain finder algorithms

Using the adjacent mouse brain slide (slide 1) used in Figure 1D, we evaluated the performance of different domain finder algorithms. For this, we manually annotated the tissue domains present in the slides analyzed using the mouse coronal P56 sample from Allen Brain Atlas^46^, which we considered as ground truth. Different algorithms (SpaGCN, Banksy, DeepST, neighborhood-based domain identification) were then used to define tissue domains, adjusting the number of domains identified to the number of domains manually annotated. Adjusted rand index (ARI) was used to compare the performance between the different algorithms. Methods with an ARI similar to the annotated domains were considered to have a better performance. In addition, domain-specific scores were obtained by calculating the maximum similarity between each manually annotated domain and the domains identified with the different algorithms using Jaccard Index.

### Segmentation-free analysis using SSAM

Segmentation-free output was produced using the SSAM package (v1.0.1) using python v3.6. In total, two rounds of analysis were performed with different parameterizations.

The first round was a SSAM de novo cell-type mapping analysis using the (x,y)-coordinates of Xenium’s provided mouse brain coronal section. SSAM was run at a resolution of 1 pixel/um and parameterized with a kernel bandwidth of 2.5. Local maximum signal points were sampled from SSAM’s vector field, and the signatures of the sampled expression vectors were normalized and clustered (by first reducing the data to 50 dimensions using PCA and applying the Louvain algorithm at a resolution of 0.3) using SSAM’s built-in de-novo functionality. The detected 91 clusters were then each assigned one out of 29 celltypes by correlating their expression centroids to the mean expression profiles of Yao et. al^17^ and correcting manually for ill-defined clusters. A SSAM cell-type map was produced filtered using a median filter to remove potential noise.

For a 3d analysis, the sample’s global gradient along the z-axis had to be corrected for the SSAM analysis to be computationally tractable. A local elevation estimator was created per molecule by building a KNN-graph of all molecules in (x,y)-space and locally propagating the molecules’ z-values in an iterative graph diffusion strategy. The local elevation estimator was subtracted from each molecule’s z-value, rendering a sample representation that was approximately geometrically centered around the z-origin at all locations. Single outliers above and below 9 μm from the new z-origin were removed. The centered spot data was subjected to a 3d-SSAM de novo analysis run at a 2.5 pixels/μm resolution, using a kernel bandwidth of 2.5 μm. Convolution edge effects were corrected for by multiplying the signal with an inverse gaussian of the same bandwidth along the z-axis, centered around the z-axis origin.

To investigate signal coherence along the z-axis, SSAM vector field was split in the middle along the z-axis, creating two 9 μm semi-sections. The correlation of the top-bottom SSAM vector field signals at each (x/y) coordinate was computed and multiplied by the 2d vector field norm at each location. Peaks in the resulting signal coherence indicator were detected in a localmax search to identify regions of interest. Out of these regions of interest (sorted by signal coherence), the regions ‘2’, ‘3’, and ‘7’ were selected for display since they covered different parts of the tissue, cell types and structures (Extended Data Figure 6A).

For the display, molecules within a 40-pixel window around a region of interest were identified and assigned a cell type by correlating the local, integrated SSAM vector field signal to expression profiled from Yao et al.^17^, similar to the earlier two-dimensional analysis. Final product was a scatter plot of the regional molecules colored by cell type, with half the region of interest removed along the x-axis to reveal the (central) spot of highest signal incoherence.

### Points2Regions

Points2Regions^32^ was used as one of the segmentation free approaches. In essence, Points2Regions is a plugin for TissUUmaps 3, intended for quick exploratory and interactive dissection of molecular patterns in *in situ* transcriptomics data. At its core, the plugin works by collecting markers in spatial bins of width *w*. Each bin thus comprises a composition of molecular markers. Adjacent bins are blurred together using a Gaussian filter parameterized by the standard deviation *σ*. Bins containing few markers are excluded based on a user-defined threshold *τ*. Bins passing the threshold are finally normalized by total count and clustered using mini-batch KMeans clustering with *k* clusters. The plugin thus comes with four tunable parameters: *w*, *σ*, *τ* and *k*, each respectively set here to 1 μm, 3 μm, 0 and 100 for all experiments.

## Author’s contributions

S.M.S and M.N. designed the study. S.M.S. obtained, formatted and described the Xenium datasets used in this study. C.M.L and S.M.S compared the Xenium datasets against other spatial technologies. S.M.S, P.C, M.C.P and K.T. annotated the cell populations. P.C. annotated the tissue domains in all datasets and S.M.S benchmarked tissue domain identification algorithms. M.D.L, L.B.K and H.R contributed to benchmarking segmentation strategies. C.W., A.A, N.I, S.T and C.A. applied segmentation-free algorithms to Xenium datasets and N.I, S.T and S.M.S evaluated the patterns identified. S.H. adapted and implemented the benchmarking pipeline for gene imputation. S.M.S, C.A and A.A contributed to data analysis while S.M.S, M.N, N.I., P.C and M.D.L. contributed to data interpretation. S.M.S., N.I, M.N, P.C, M.D.L, C.A, C.W and A.A. contributed to writing the manuscript. All authors read and approved the content of this manuscript.

## Data availability

The Xenium datasets used in this analysis were provided by 10X Genomics. The three mouse brain coronal sections (“ms brain multisection”) are publically available datasets and can be downloaded at from the 10X website (https://www.10xgenomics.com/resources/datasets). The mouse brain full coronal and half coronal sections (named as “ms brain coronal” and “ms brain ROI” in Figure 1B), as well as the human breast sections are available upon request.

## Code availability

All the code used in this analysis can be found in the following github repository: https://github.com/Moldia/Xenium_benchmarking. Since different tools require their own environment, analysis are divided in folders, providing different Conda recipes files to recreate the environments needed to reproduce the analysis.

## Acknowledgements

The datasets generated using Xenium and described in this manuscript have been provided by 10x Genomics, Inc. and correspond to data generated using an in-development gene panel, chemistry and instrument. Thus, they may not be indicative of final application performance.

## References

1. Marx V. Method of the Year: spatially resolved transcriptomics. Nat Methods. 2021;18(1):9–14. doi:10.1038/S41592-020-01033-Y

2. Moses L, Pachter L. Museum of spatial transcriptomics. doi:10.1038/s41592-022-01409-2

3. Ståhl PL, Salmén F, Vickovic S, et al. Visualization and analysis of gene expression in tissue sections by spatial transcriptomics. Science (80-). 2016;353(6294):78–82. doi:10.1126/SCIENCE.AAF2403/SUPPL_FILE/AAF2403_STAHL_SM.PDF

4. Rodriques SG, Stickels RR, Goeva A, et al. Slide-seq: A scalable technology for measuring genome-wide expression at high spatial resolution. Science (80-). 2019;363(6434):1463–1467. doi:10.1126/SCIENCE.AAW1219/SUPPL_FILE/AAW1219S1.MOV

5. Chen A, Liao S, Cheng M, et al. Spatiotemporal transcriptomic atlas of mouse organogenesis using DNA nanoball-patterned arrays. Cell. 2022; 185(10):1777–1792.e21. doi:10.1016/J.CELL.2022.04.003

6. Chen KH, Boettiger AN, Moffitt JR, Wang S, Zhuang X. Spatially resolved, highly multiplexed RNA profiling in single cells. Science (80-). 2015;348(6233). doi:10.1126/SCIENCE.AAA6090/SUPPL_FILE/CHEN-SM.PDF

7. Shah S, Lubeck E, Zhou W, Cai L. In Situ Transcription Profiling of Single Cells Reveals Spatial Organization of Cells in the Mouse Hippocampus. Neuron. 2016;92(2):342–357. doi:10.1016/J.NEURON.2016.10.001

8. Borm LE, Mossi Albiach A, Mannens CCA, et al. Scalable in situ single-cell profiling by electrophoretic capture of mRNA using EEL FISH. Nat Biotechnol 2022. Published online September 22, 2022:1–10. doi:10.1038/s41587-022-01455-3

9. Gyllborg D, Langseth CM, Qian X, et al. Hybridization-based in situ sequencing (HybISS) for spatially resolved transcriptomics in human and mouse brain tissue. Nucleic Acids Res. 2020;48(19):e112. doi:10.1093/nar/gkaa792

10. Biancalani T, Scalia G, Buffoni L, et al. Deep learning and alignment of spatially resolved single-cell transcriptomes with Tangram. Nat Methods 2021 1811. 2021;18(11):1352–1362. doi:10.1038/s41592-021-01264-7

11. Sountoulidis A, Liontos A, Nguyen HP, et al. SCRINSHOT enables spatial mapping of cell states in tissue sections with single-cell resolution. PLOS Biol. 2020;18(11):e3000675. doi:10.1371/JOURNAL.PBIO.3000675

12. Janesick A, Shelansky R, Gottscho AD, et al. High resolution mapping of the breast cancer tumor microenvironment using integrated single cell, spatial and in situ analysis of FFPE tissue. bioRxiv. Published online November 3, 2022:2022.10.06.510405. doi:10.1101/2022.10.06.510405

13. Tasic B, Yao Z, Graybuck LT, et al. Shared and distinct transcriptomic cell types across neocortical areas. Nat 2018 5637729. 2018;563(7729):72–78. doi:10.1038/s41586-018-0654-5

14. Zeisel A, Hochgerner H, Lönnerberg P, et al. Molecular Architecture of the Mouse Nervous System. Cell. 2018;174(4):999–1014.e22. doi:10.1016/j.cell.2018.06.021

15. Harris KD, Hochgerner H, Skene NG, et al. Classes and continua of hippocampal CA1 inhibitory neurons revealed by single-cell transcriptomics. PLOS Biol. 2018;16(6):e2006387. doi:10.1371/JOURNAL.PBIO.2006387

16. Hodge RD, Bakken TE, Miller JA, et al. Conserved cell types with divergent features in human versus mouse cortex. Nat 2019 5737772. 2019;573(7772):61–68. doi:10.1038/s41586-019-1506-7

17. Yao Z, van Velthoven CTJ, Nguyen TN, et al. A taxonomy of transcriptomic cell types across the isocortex and hippocampal formation. Cell. 2021;184(12): 3222–3241.e26. doi:10.1016/J.CELL.2021.04.021

18. Codeluppi S, Borm LE, Zeisel A, et al. Spatial organization of the somatosensory cortex revealed by osmFISH. Nat Methods 2018 1511. 2018;15(11):932–935. doi:10.1038/s41592-018-0175-z

19. Qian X, Harris KD, Hauling T, et al. Probabilistic cell typing enables fine mapping of closely related cell types in situ. Nat Methods. 2020;17(1):101–106. doi:10.1038/s41592-019-0631-4

20. Maniatis S, Petrescu J, Phatnani H. Spatially resolved transcriptomics and its applications in cancer. Curr Opin Genet Dev. 2021;66:70–77. doi:10.1016/J.GDE.2020.12.002

21. Zhang M, Eichhorn SW, Zingg B, et al. Spatially resolved cell atlas of the mouse primary motor cortex by MERFISH. Nat 2021 5987879. 2021;598(7879):137–143. doi:10.1038/s41586-021-03705-x

22. Alon S, Goodwin DR, Sinha A, et al. Expansion sequencing: Spatially precise in situ transcriptomics in intact biological systems. Science (80-). 2021;371(6528). doi:10.1126/SCIENCE.AAX2656/SUPPL_FILE/AAX2656_TABLESS1-S6ANDS9-S14.XLSX

23. Shi H, He Y, Zhou Y, et al. Spatial Atlas of the Mouse Central Nervous System at Molecular Resolution. bioRxiv. Published online June 22, 2022:2022.06.20.496914. doi:10.1101/2022.06.20.496914

24. Lee H, Marco Salas S, Gyllborg D, Nilsson M. Direct RNA targeted in situ sequencing for transcriptomic profiling in tissue. Sci Reports 2022 121. 2022;12(1):1–9. doi:10.1038/s41598-022-11534-9

25. Zhang Y, Miller JA, Park J, et al. Reference-based cell type matching of spatial transcriptomics data. bioRxiv. Published online May 12, 2022:2022.03.28.486139. doi:10.1101/2022.03.28.486139

26. Chen X, Sun YC, Church GM, Lee JH, Zador AM. Efficient in situ barcode sequencing using padlock probe-based BaristaSeq. Nucleic Acids Res. 2018;46(4). doi:10.1093/nar/gkx1206

27. Beucher S, Lantuéjoul C. Use of Watersheds in Contour Detection. Vol 132.; 1979.

28. Stringer C, Wang T, Michaelos M, Pachitariu M. Cellpose: a generalist algorithm for cellular segmentation. Nat Methods 2020 181. 2020;18(1):100–106. doi:10.1038/s41592-020-01018-x

29. Petukhov V, Xu RJ, Soldatov RA, et al. Cell segmentation in imaging-based spatial transcriptomics. Nat Biotechnol 2021 403. 2021;40(3):345–354. doi:10.1038/s41587-021-01044-w

30. He Y, Tang X, Huang J, et al. ClusterMap for multi-scale clustering analysis of spatial gene expression. Nat Commun 2021 121. 2021;12(1):1–13. doi:10.1038/s41467-021-26044-x

31. Park J, Choi W, Tiesmeyer S, et al. Cell segmentation-free inference of cell types from in situ transcriptomics data. Nat Commun 2021 121. 2021;12(1):1–13. doi:10.1038/s41467-021-23807-4

32. Andersson A, Behanova A, Avenel C, Wählby C, Malmberg F. POINTS2REGIONS: FAST INTERACTIVE CLUSTERING OF IN SITU TRANSCRIPTOMICS DATA. doi:10.1101/2022.12.07.519086

33. Li B, Zhang W, Guo C, et al. Benchmarking spatial and single-cell transcriptomics integration methods for transcript distribution prediction and cell type deconvolution. Nat Methods 2022 196. 2022; 19(6):662–670. doi:10.1038/s41592-022-01480-9

34. Lopez R, Nazaret A, Langevin M, et al. A joint model of unpaired data from scRNA-seq and spatial transcriptomics for imputing missing gene expression measurements. Published online May 6, 2019. doi:10.48550/arxiv.1905.02269

35. Abdelaal T, Mourragui S, Mahfouz A, Reinders MJT. SpaGE: Spatial Gene Enhancement using scRNA-seq. Nucleic Acids Res. 2020;48(18):E107–E107. doi:10.1093/nar/gkaa740

36. Cang Z, Nie Q. Inferring spatial and signaling relationships between cells from single cell transcriptomic data. Nat Commun 2020 111. 2020;11(1):1–13. doi:10.1038/s41467-020-15968-5

37. Moriel N, Senel E, Friedman N, Rajewsky N, Karaiskos N, Nitzan M. NovoSpaRc: flexible spatial reconstruction of single-cell gene expression with optimal transport. Nat Protoc 2021 169. 2021;16(9):4177–4200. doi:10.1038/s41596-021-00573-7

38. Palla G, Spitzer H, Klein M, et al. Squidpy: a scalable framework for spatial omics analysis. Nat Methods 2022 192. 2022;19(2):171–178. doi:10.1038/s41592-021-01358-2

39. Dries R, Zhu Q, Dong R, et al. Giotto: a toolbox for integrative analysis and visualization of spatial expression data. Genome Biol. 2021;22(1):1–31. doi:10.1186/S13059-021-02286-2/FIGURES/6

40. DeTomaso D, Yosef N. Hotspot identifies informative gene modules across modalities of single-cell genomics. Cell Syst. 2021;12(5):446–456.e9. doi:10.1016/J.CELS.2021.04.005

41. Hao M, Hua K, Zhang X. SOMDE: A scalable method for identifying spatially variable genes with self-organizing map. Bioinformatics. 2021;37(23):4392–4398. doi:10.1093/BIOINFORMATICS/BTAB471

42. Svensson V, Teichmann SA, Stegle O. SpatialDE: identification of spatially variable genes. Nat Methods 2018 155. 2018;15(5):343–346. doi:10.1038/nmeth.4636

43. Ruiz-Moreno C, Salas SM, Samuelsson E, et al. Harmonized single-cell landscape, intercellular crosstalk and tumor architecture of glioblastoma. bioRxiv. Published online August 27, 2022:2022.08.27.505439. doi:10.1101/2022.08.27.505439

44. Singhal V, Chou N, Lee J, et al. BANKSY: A Spatial Omics Algorithm that Unifies Cell Type Clustering and Tissue Domain Segmentation. bioRxiv. Published online April 15, 2022:2022.04.14.488259. doi:10.1101/2022.04.14.488259

45. Xu C, Jin X, Wei S, et al. DeepST: identifying spatial domains in spatial transcriptomics by deep learning. Nucleic Acids Res. Published online October 17, 2022. doi:10.1093/NAR/GKAC901

46. Wang Q, Ding SL, Li Y, et al. The Allen Mouse Brain Common Coordinate Framework: A 3D Reference Atlas. Cell. 2020;181(4):936–953.e20. doi:10.1016/J.CELL.2020.04.007

47. Mah CK, Ahmed N, Lam D, et al. Bento: A toolkit for subcellular analysis of spatial transcriptomics data. bioRxiv. Published online June 13, 2022:2022.06.10.495510. doi:10.1101/2022.06.10.495510

48. Hu J, Li X, Coleman K, et al. SpaGCN: Integrating gene expression, spatial location and histology to identify spatial domains and spatially variable genes by graph convolutional network. Nat Methods 2021 1811. 2021;18(11):1342–1351. doi:10.1038/s41592-021-01255-8

