## Extended Data Figures for "Optimizing Xenium In Situ data utility by quality assessment and best practice analysis workflows"

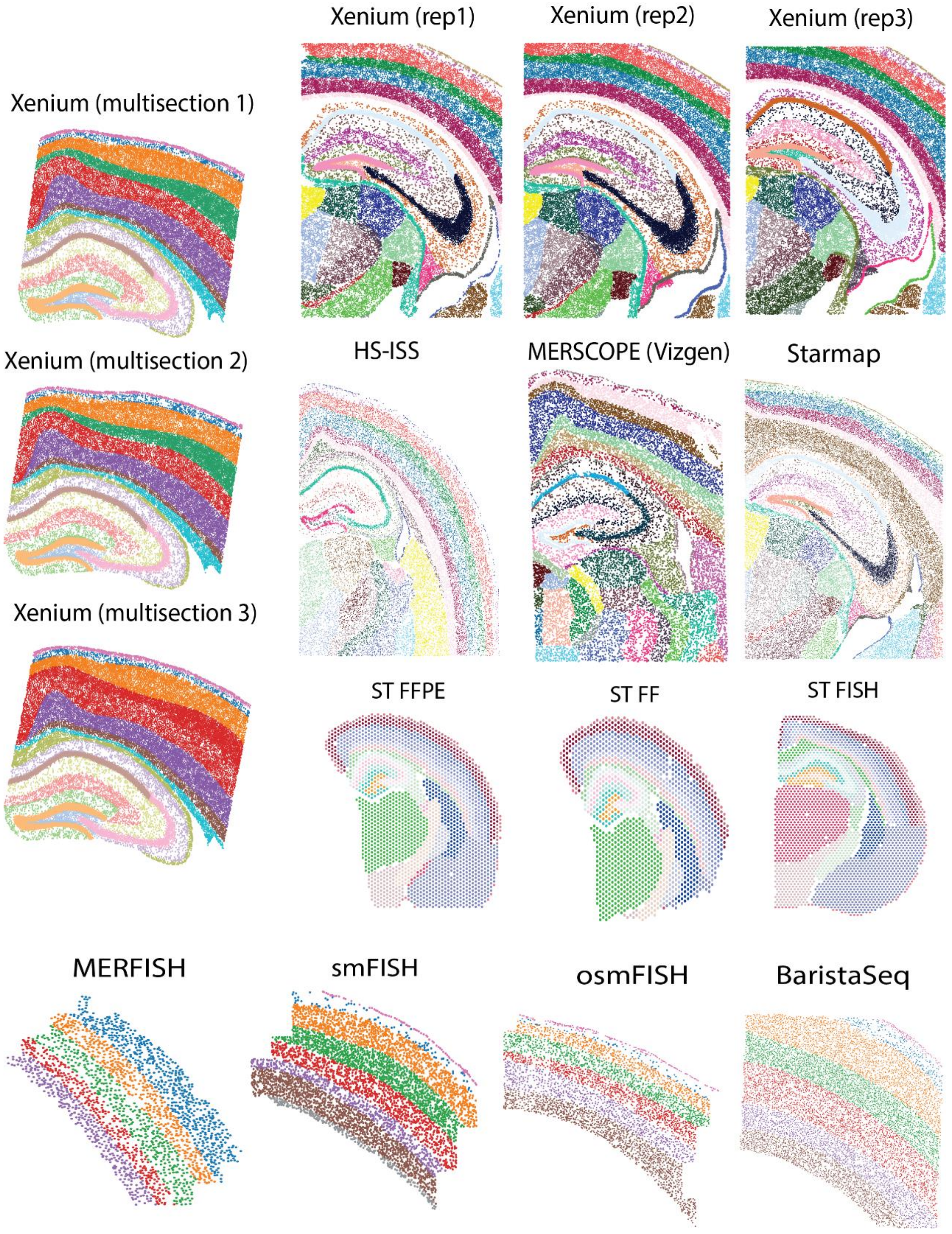


**Extended Data figure 1: Spatial maps of the mouse brain sections used to compare Xenium to other technologies.** Individual cells or spatial transcriptomics/Visium spots are colored depending on the annotated region they belong to.

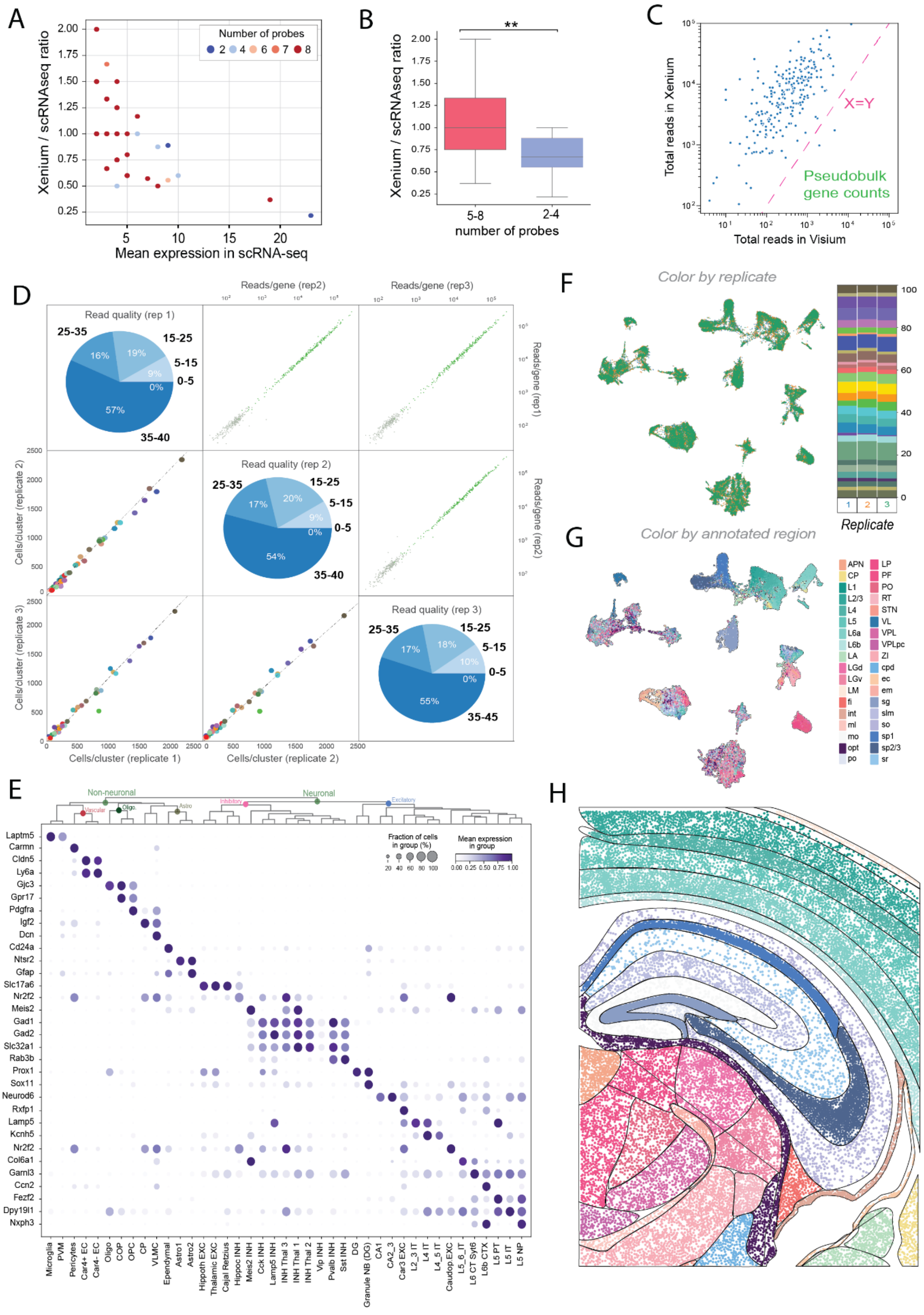

A
Number of probes
2
4
6
7
8
Xenium / scRNAseq ratio
Mean expression in scRNA-seq
B
**
Xenium / scRNAseq ratio
5-8
2-4
number of probes
C
Total reads in Xenium
Total reads in Visium
X=Y
Pseudobulk gene counts
D
Read quality (rep 1)
25-35
15-25
5-15
0-5
35-40
16%
19%
9%
0%
57%
Reads/gene (rep2)
Reads/gene (rep3)
Reads/gene (rep1)
Read quality (rep 2)
25-35
15-25
5-15
0-5
35-40
17%
20%
9%
0%
54%
Reads/gene (rep2)
Cells/cluster (replicate 2)
Read quality (rep 3)
25-35
15-25
5-15
0-5
35-45
17%
18%
10%
0%
55%
Cells/cluster (replicate 3)
Cells/cluster (replicate 1)
Cells/cluster (replicate 2)
E
Non-neuronal
Neuronal
Vascular
Oligo
Astro
Inhibitory
Excitatory
Fraction of cells in group (%)
Mean expression in group
Laptm5
Carmn
Cldn5
Ly6a
Gjc3
Gpr17
Pdgfra
Igf2
Dcn
Cd24a
Ntsr2
Gfap
Slc17a6
Nr2f2
Meis2
Gad1
Gad2
Slc32a1
Rab3b
Prox1
Sox11
Neurod6
Rxfp1
Lamp5
Kcnh5
Nr2f2
Col6a1
Garnl3
Ccn2
Fezf2
Dpy19l1
Nxph3
Microglia
PVM
Pericytes
Car4+ EC
Car4- EC
Oligo
COP
OPC
CP
VLMC
Ependymal
Astro1
Astro2
Hippoth EXC
Thalamic EXC
Cajal Retzius
Hippoc INH
Meis2 INH
Cck INH
Lamp5 INH
INH Thal 3
INH Thal 1
INH Thal 2
Vip INH
Pvalb INH
Sst INH
DG
Granule NB (DG)
CA1
CA2_3
Car3 EXC
L2_3 IT
L4 IT
L4_5 IT
Caudop.EXC
L5_6_IT
L6 CT Syt6
L6b CTX
L5 PT
L5 IT
L5 NP
F
Color by replicate
Replicate
G
Color by annotated region
APN
CP
L1
L2/3
L4
L5
L6a
L6b
LA
LGd
LGv
LM
fi
int
ml
mo
opt
po
LP
PF
PO
RT
STN
VL
VPL
VPLpc
ZI
cpd
ec
em
sg
slm
so
sp1
sp2/3
sr
H

**Extended Data figure 2: Complementary information regarding cell typing in the mouse brain sections and comparisons with other SRT technologies. A.** Scatter plot representing the correlation (r=0.5) between the mean expression of each gene in scRNAseq compared with its Xenium/scRNA-seq ratio for genes detected in both datasets. **B.** Boxplot of the Xenium/scRNA-seq efficiency ratios when grouping the genes by the number of padlock probes designed against them. Boxplots represent the distribution of the efficiencies, divided in quartiles, where the central line represents the median efficiency. T-test was performed to compare the groups and significance is encoded as follows: p value < 0.01 **C.** Scatter plot of the reads captured in Visium compared to the reads captured by Xenium for each specific read in the same cortical region of the mouse brain. **D.** Composite representation of the consistency between technical replicates. Consistency in gene counts (upper triangular matrix) and cluster counts (lower triangular matrix) is shown using scatter plots between the three replicates. In the upper triangular matrix, transcripts in the panel are represented in green, while negative control barcodes are represented in gray.Cell type colors shown in the lower triangular matrix correspond to the ones used in Figure 1E. Diagonal pie plots represent the phred-score based read quality in the three replicates. **E.** Dotplot representing the expression pattern of the top differentially expressed gene for every cell type defined in Figure 1C across all cell types. **F.** UMAP representation of cells found in the 3 adjacent mouse brain slides colored by slide (“replicate”). **G.** UMAP representation of cells found in the 3 adjacent mouse brain slides colored by manually annotated regions. **H.** Spatial map of cells identified in mouse brain region of interest (replicate 1) colored by regions defined in Extended Data Figure 2D. Annotated domains are plotted using the outline of each region.

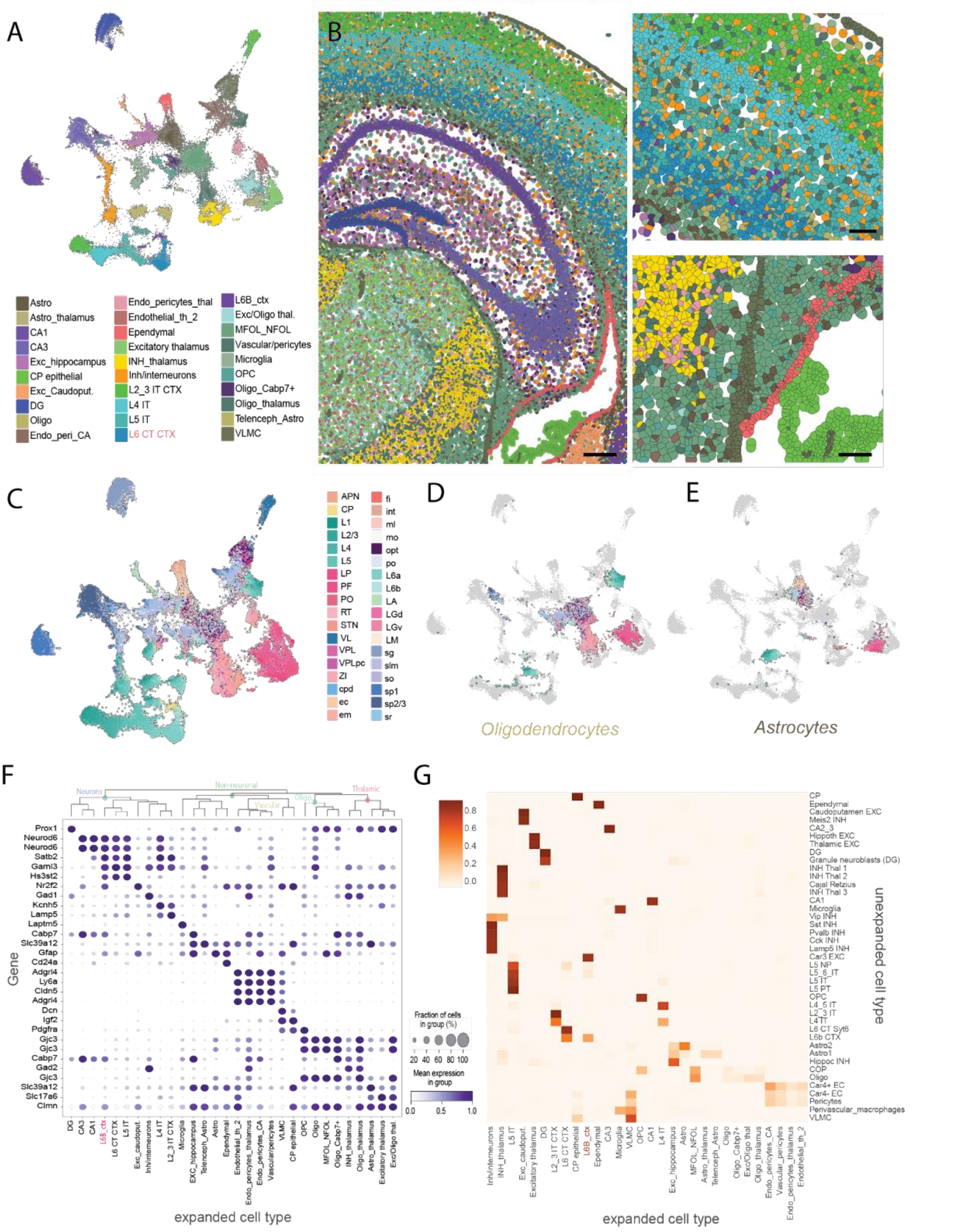


**Extended Data figure 3: Analysis of expanded cells in the mouse brain region. A.** UMAP representation of clusters by Xenium in the 3 adjacent mouse brain regions using expanded cells. **B.** Spatial map of the clusters identified in panel A in replicate 1 (right) and two regions of interest in the cortex (left, up) and the ependymal / CP (left, down). Cells are represented using the cell boundaries identified by Xenium's segmentation algorithm after the expansion. Colors correspond to clusters identified in panel A. **C-E.** UMAP representation of expanded cells found in the 3 adjacent mouse brain slides colored by manually annotated regions including all cells (C), oligodendrocytes (D) and astrocytes (E). **F.** Dotplot representing the expression pattern of the top differentially expressed gene for every cluster defined in Extended Data Figure 3A across all cell types. **G.** Heatmap representing the correspondence between cell types identified in unexpanded and expanded cells.

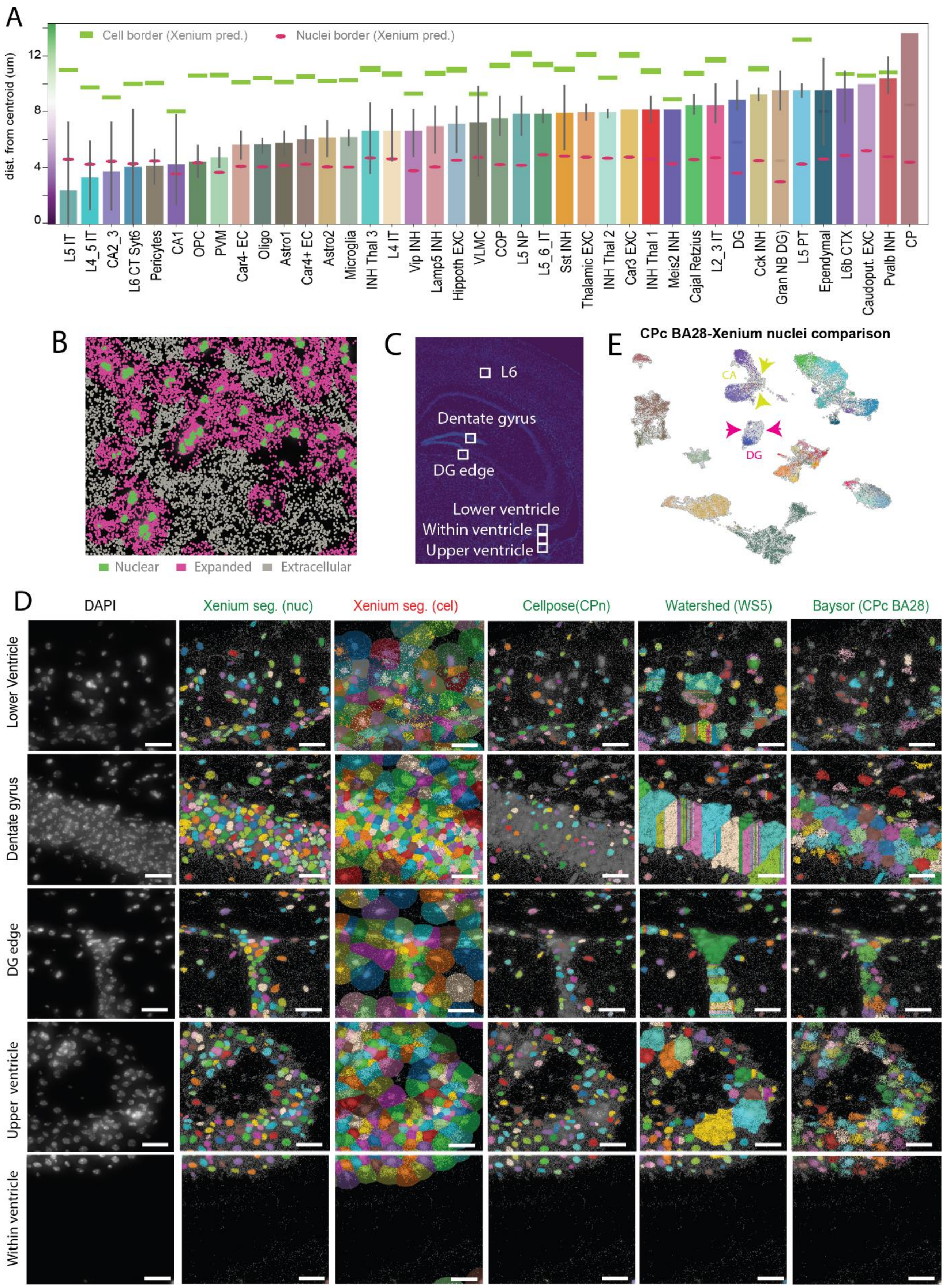


**Extended Data figure 4: A.**Bar plot indicating the distance to the nuclei at which the correlation to the domain-specific background signature becomes higher than to the nuclei cell type and domain-specific expression signature for all of the clusters identified at Figure 1C with at least 5000 reads assigned to the assessed domain, same as in Figure 2C. **B.** Expression map representing the location of decoded reads in a region of interest selected from the sample named mouse brain replicate 1. Reads are overlaid on top of their corresponding nuclei staining (DAPI) and colored depending on their assignment when segmenting cells in (1) nuclear, if it was detected inside segmented cells before the expansion, (2) cytoplasmic, if it was detected inside segmented cells after the expansion, or (3) extracellular, if it wasn't assigned to any cell. **C** Localization of regions of interest represented in Extended Data Figure 4E and Figure 2D. **D.** Regions of interest representing the cells identified using different segmentation algorithms in a region of interest outlined in Extended Data 4C. DAPI background is represented as a background and individual isolated color-specific masks represent individual cells. Segmentation strategies were selected to represent different segmentation outputs, as described in Figure 2D. Scale bars represent 25 μm. **E.** UMAP representing the cells segmented by Xenium's algorithm (gray) when clustered together with Cellpose followed by Baysor segmentation (CPc BA28, no expansion). These cells are colored by their identified cell type, following the coloring scheme described in Figure 1C. Arrows highlight differences between segmentation methods in the dentate gyrus (pink) and the hippocampus (CA).

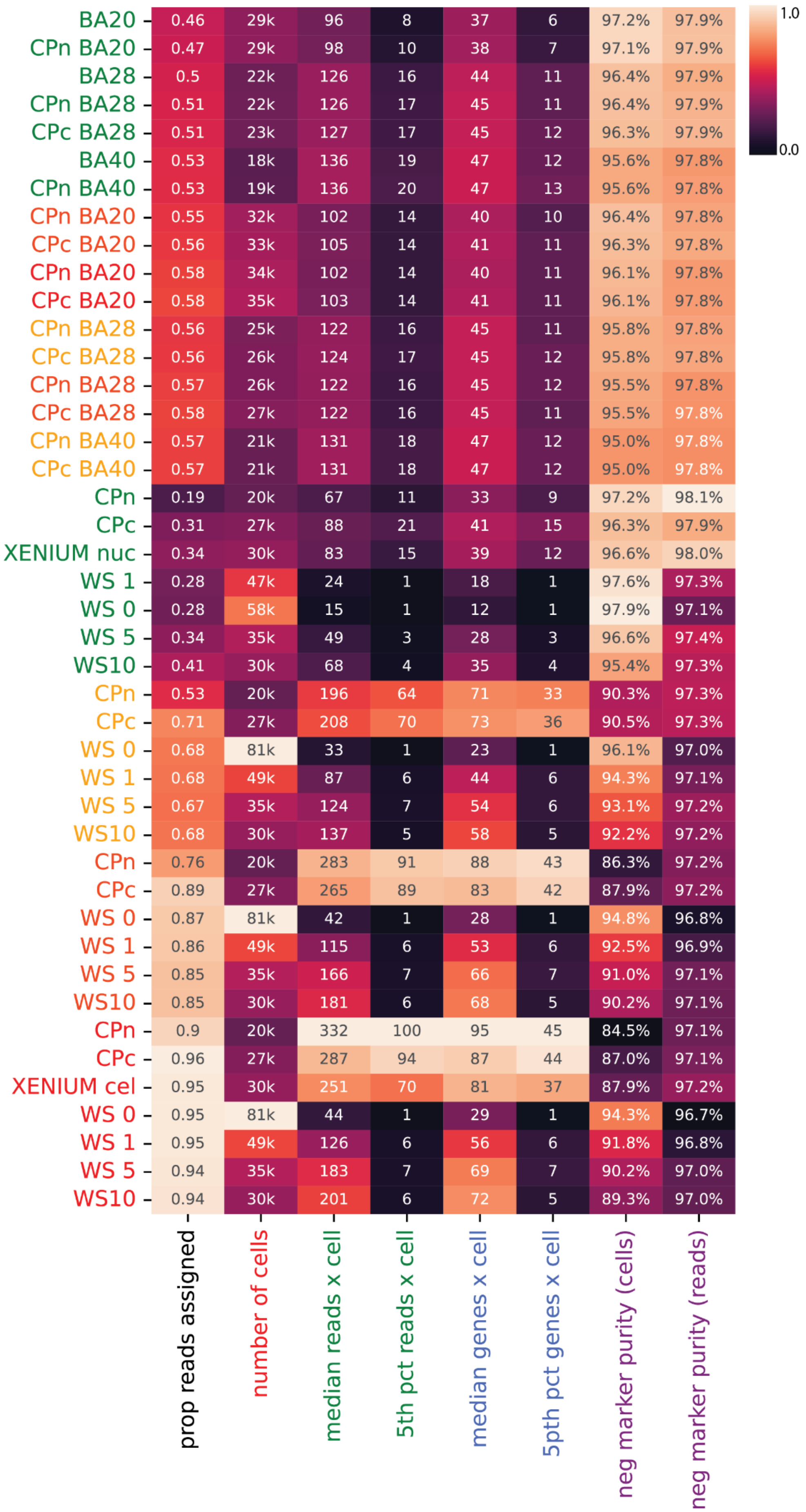
BA20 0.46 29k 96 8 37 6 97.2% 97.9%
CPn BA20 0.47 29k 98 10 38 7 97.1% 97.9%
BA28 0.5 22k 126 16 44 11 96.4% 97.9%
CPn BA28 0.51 22k 126 17 45 11 96.4% 97.9%
CPc BA28 0.51 23k 127 17 45 12 96.3% 97.9%
BA40 0.53 18k 136 19 47 12 95.6% 97.8%
CPn BA40 0.53 19k 136 20 47 13 95.6% 97.8%
CPn BA20 0.55 32k 102 14 40 10 96.4% 97.8%
CPc BA20 0.56 33k 105 14 41 11 96.3% 97.8%
CPn BA20 0.58 34k 102 14 40 11 96.1% 97.8%
CPc BA20 0.58 35k 103 14 41 11 96.1% 97.8%
CPn BA28 0.56 25k 122 16 45 11 95.8% 97.8%
CPc BA28 0.56 26k 124 17 45 12 95.8% 97.8%
CPn BA28 0.57 26k 122 16 45 12 95.5% 97.8%
CPc BA28 0.58 27k 122 16 45 11 95.5% 97.8%
CPn BA40 0.57 21k 131 18 47 12 95.0% 97.8%
CPc BA40 0.57 21k 131 18 47 12 95.0% 97.8%
CPn 0.19 20k 67 11 33 9 97.2% 98.1%
CPc 0.31 27k 88 21 41 15 96.3% 97.9%
XENIUM nuc 0.34 30k 83 15 39 12 96.6% 98.0%
WS 1 0.28 47k 24 1 18 1 97.6% 97.3%
WS 0 0.28 58k 15 1 12 1 97.9% 97.1%
WS 5 0.34 35k 49 3 28 3 96.6% 97.4%
WS10 0.41 30k 68 4 35 4 95.4% 97.3%
CPn 0.53 20k 196 64 71 33 90.3% 97.3%
CPc 0.71 27k 208 70 73 36 90.5% 97.3%
WS 0 0.68 81k 33 1 23 1 96.1% 97.0%
WS 1 0.68 49k 87 6 44 6 94.3% 97.1%
WS 5 0.67 35k 124 7 54 6 93.1% 97.2%
WS10 0.68 30k 137 5 58 5 92.2% 97.2%
CPn 0.76 20k 283 91 88 43 86.3% 97.2%
CPc 0.89 27k 265 89 83 42 87.9% 97.2%
WS 0 0.87 81k 42 1 28 1 94.8% 96.8%
WS 1 0.86 49k 115 6 53 6 92.5% 96.9%
WS 5 0.85 35k 166 7 66 7 91.0% 97.1%
WS10 0.85 30k 181 6 68 5 90.2% 97.1%
CPn 0.9 20k 332 100 95 45 84.5% 97.1%
CPc 0.96 27k 287 94 87 44 87.0% 97.1%
XENIUM cel 0.95 30k 251 70 81 37 87.9% 97.2%
WS 0 0.95 81k 44 1 29 1 94.3% 96.7%
WS 1 0.95 49k 126 6 56 6 91.8% 96.8%
WS 5 0.94 35k 183 7 69 7 90.2% 97.0%
WS10 0.94 30k 201 6 72 5 89.3% 97.0%
prop reads assigned
number of cells
median reads x cell
5th pct reads x cell
median genes x cell
5pth pct genes x cell
neg marker purity (cells)
neg marker purity (reads)
1.0
0.0

**Extended Data Figure 5: Performance of segmentation algorithms.** Heatmap representing the performance of different segmentation algorithms and hyperparameters (y-axis) using different quality metrics (x-axis). Segmentation methods included Cellpose (CPn: nuclei, CPc: cyto models),binning (bins), clustermap (clustermap), watershed (WA), Baysor (BA) and cellpose combined with Baysor (CPc BA/CPn BA). Xenium segmentation were also included in the comparison (XENIUM cel, XENIUM nuc). Hyperparameters for each method are described in methods. Methods on the y-axis were colored depending on the expansion performed after segmentation (green: 0 um, yellow: 5.3 μm, orange: 10.6 μm, red: 15.9 μm)

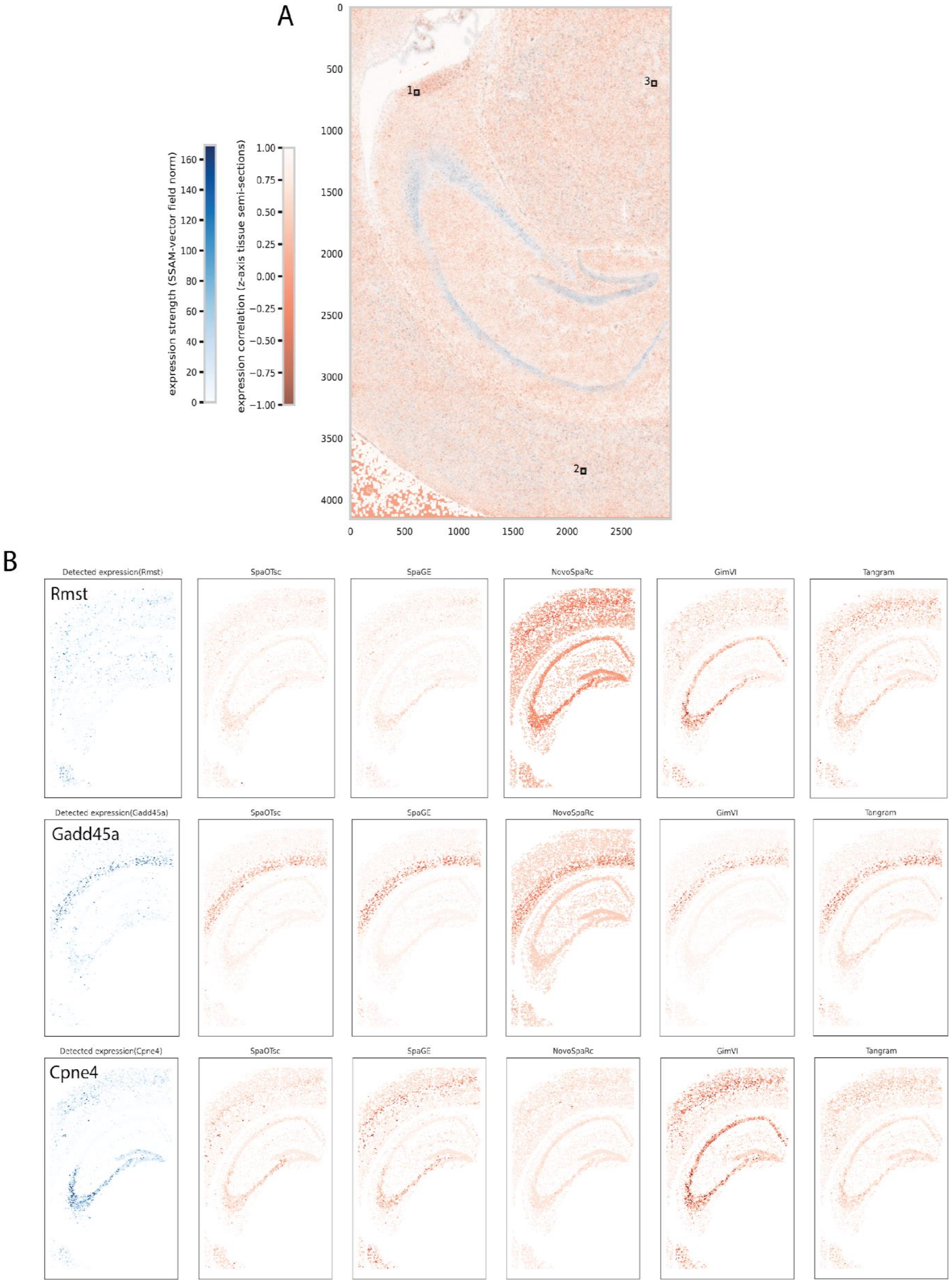
A
expression strength (SSAM-vector field norm)
expression correlation (z-axis tissue semi-sections)
1
2
3
B
Detected expression(Rmst)
Rmst
SpaOTsc
SpaGE
NovoSpaRc
GimVI
Tangram
Detected expression(Gadd45a)
Gadd45a
Detected expression(Cpne4)
Cpne4

**Extended Data Figure 6. Differences in 3-D signatures and expression profiles of imputed genes**. **A.** Spatial map of the mouse brain section used for SSAM analysis. The expression strength (blue) and expression correlation in the Z-axis (red) are represented. Regions with a high expression strength and low expression correlation of the Z-axis indicate the presence of overlapping cells. **B.**Spatial maps representing the detected expression of different genes (columns) and their corresponding imputed expression profiles (rows) using the benchmarked algorithms from Figure 4A. Genes that presented a high (Gadd45a) , medium (Cpne4) and low (Rmst) imputation accuracy score are represented to highlight the gene-specific variability of gene imputation methods.
